# Crosstalk of platelets with macrophages and fibroblasts aggravates inflammation, aortic wall stiffening and osteopontin release in abdominal aortic aneurysm

**DOI:** 10.1101/2022.12.01.518674

**Authors:** MU Wagenhäuser, J Mulorz, KJ Krott, A Bosbach, T Feige, YH Rhee, M Chatterjee, N Petzold, C Böddeker, W Ibing, I Krüger, AM Popovic, A Roseman, JM Spin, PS Tsao, H Schelzig, M Elvers

**Affiliations:** Department of Vascular- and Endovascular Surgery, University Hospital Düsseldorf, Heinrich-Heine University, Düsseldorf, Germany; Department of Pharmacology, Experimental Therapy and Toxicology, University Hospital Tübingen, Tübingen, Germany; VA Palo Alto Health Care System, Palo Alto, CA, United States; Stanford University, Department of Cardiovascular Medicine, Stanford, CA, United States

**Keywords:** platelets, abdominal aortic aneurysm, inflammation, ECM remodeling, osteopontin

## Abstract

Abdominal aortic aneurysm (AAA) is a highly lethal disease with progressive dilatation of the abdominal aorta accompanied by degradation and remodelling of the vessel wall due to chronic inflammation. Platelets play an important role in cardiovascular diseases but their role in AAA is poorly understood. The present study revealed that platelets play a crucial role in promoting AAA through modulation of inflammation and degradation of the ECM. They are responsible for the up-regulation of *SPP1* (*osteopontin, OPN)* gene expression in macrophages and aortic tissue, which triggers inflammation and remodeling but also platelet adhesion and migration into the abdominal aortic wall and the intraluminal thrombus (ILT). Further, enhanced platelet activation and pro-coagulant activity results in elevated gene expression of various cytokines, *Mmp9* and *Col1a1* in macrophages and *Il-6* and *Mmp9* in fibroblasts. Enhanced platelet activation and pro-coagulant activity was also detected in AAA patients. Further, we detected platelets and OPN in the vessel wall and in the ILT of patients who underwent open repair of AAA. Platelet depletion in experimental murine AAA reduced inflammation and ECM remodeling, with reduced elastin fragmentation and aortic diameter expansion. Of note, OPN co-localized with platelets, suggesting a potential role of OPN for the recruitment of platelets into the ILT and the aortic wall. In conclusion, our data strongly supports the potential relevance of anti-platelet therapy to reduce AAA progression and rupture in AAA patients.

**Translational perspective:** Abdominal aortic aneurysm (AAA) is a severe cardiovascular disease (CVD) with high mortality. Since the role of platelets is unclear, we explored platelet-mediated processes in the pathogenesis of AAA. Results from platelet depleted mice and patients with AAA revealed that platelets modulate inflammatory and stiffness-related gene expression of macrophages and fibroblasts. Further, platelets induce the release of osteopontin important for the recruitment of platelets to the aortic wall and to the intraluminal thrombus (ILT). Consequently, platelet depletion significantly reduced aneurysm growth. Thus, therapeutic targeting of platelet activation might be crucial for the treatment of patients to reduce AAA formation and progression.

## 1. Introduction

Abdominal aortic aneurysm (AAA) is a localized, progressive dilatation of the abdominal aorta caused by chronic inflammation resulting in adverse remodeling of the aortic wall. Well-established risk factors include age, hypertension, male gender, and smoking. In Western countries, the incidence of AAA is ∼0.4–0.67% annually and reaches up to 5–10% in men over age 65^1^. As AAA diameter increases, the risk of fatal rupture rises exponentially. Despite remarkable advances in understanding the underlying pathological mechanisms during the past years, endovascular and open repair remain the only AAA treatment options. Although potential candidates exist, such as metformin, non-invasive drug-based therapies are pending clinical implementation, since prospective randomized trials have either not demonstrated efficacy or are yet to be performed^2–6^.

Characteristic features of AAA pathophysiology are the recruitment of infiltrating inflammatory leukocytes, together with the activation of proteolytic enzymes leading to extracellular matrix degradation, which weakens the aortic vessel wall^7,8^. In this context the formation of an intraluminal thrombus (ILT) may be critical for the progress of the disease, since the ILT modulates both, biomechanical parameters and inflammatory processes in the AAA vessel wall^9,10^. The vast majority of AAAs are accompanied by an ILT, although its disease-specific role is still widely debated^11,12^. The key elements of inflammatory processes and the formation of an ILT suggest that platelets and their activation might play a significant role for AAA initiation and progression, particularly as evidence has accumulated for platelet-based acceleration of various cardiovascular diseases^13^.

Various preclinical animal models exist, allowing mechanistic insights into various aspects of AAA disease^14^. Of these, the angiotensin-II (Ang-II) infusion model provides features of dissection and aneurysm formation, and has demonstrated a beneficial effect in AAA rupture rates and reduced macrophage recruitment into the AAA wall in response to antiplatelet drugs^15^. Further, the platelet integrin α_IIb_β_3_ inhibitor abciximab and/or a P2Y_12_ receptor antagonist both blocked AAA growth in a rat xenograft aneurysm model^16,17^. To date, no study has explored the role of platelets in the pancreatic porcine elastase (PPE) infusion model. The PPE model reproduces many features of human AAA and was therefore used in this study^14^. Small clinical trials have supported the hypothesis that platelet activation contributes to AAA development and progression, because antiplatelet medication with low-dose aspirin has been shown to be beneficial for patients with small AAAs, as indicated by decreased aneurysm size and less frequent need for surgical repair^18^. In patients with large AAAs, platelet activation is enhanced with increased levels of soluble CD40L, TAT complex formation, P-selectin exposure, and fibrinogen binding^19^.

In conclusion, both experimental and clinical data strongly suggest that, in AAA, impaired endothelial function causes platelet activation, leading to ILT formation and promoting a persistent renewal of cellular activity at the luminal interface via aggregation, entrapment, and recruitment of activated platelets, establishing a chronic inflammatory response. Thus, platelet activation may play a pivotal role in AAA formation and progression. Despite these theoretical considerations and encouraging results in preclinical animal models and small clinical trials, there is a significant lack of understanding on how platelets contribute to AAA development and progression. More insights into the specific role of platelets in AAA disease and their activation mechanisms are urgently needed to allow the development of appropriate antiplatelet targets. Here, we present findings regarding platelet contribution to AAA disease opening the way for potential novel therapies.

## 2. Methods

### 2.1 Study approval

All animal experiments were conducted according to the Declaration of Helsinki and German respectively U.S. law for the welfare of animals. The protocol was reviewed and approved by the local government authority. Germany: Heinrich-Heine-University Animal Care Committee and by the State Agency for Nature, Environment and Consumer Protection of North Rhine-Westphalia LANUV, Recklinghausen, NRW, Germany Permit Numbers 81-02.04.2018.A409 and 81-02.05.40.21.041. U.S.: C57BL/6 wild-type male mice were purchased from the Jackson Laboratory. Animals were housed in a temperature-controlled and humidity-controlled room under a 12-h light/dark cycle (6:30 am/6:30 pm). All animal protocols were approved by the VA Institutional Animal Care and Use Committee and followed the National Institutes of Health and U.S. Department of Agriculture Guidelines for Care and Use of Animals in Research.

Experiments with human tissues and blood were reviewed and approved by the Ethics Committee of the Heinrich-Heine-University, who approved the collection and analysis of the tissue samples. Subjects provided informed consent prior to their participation in the study (patients’ consent): Permitted ethical votes; ID: 2018-140-KFogU, 5731R (2018-222_1) and 2018-248-FmB. The study was conducted in accordance with Declaration of Helsinki principles and the International Council for Harmonization Guidelines on Good Clinical Practice.

### 2.2 Human AAA samples

Fresh citrate-anticoagulated blood was collected from healthy volunteers between 18 and 70 years of age into Vacutainer^®^ sodium citrate tubes (BD Vacutainer^®^, #367714). Blood of older (>60 years) healthy volunteers was used as age-matched controls (AMCs) for the comparison with AAA patients (study number: 2018-140-KFogU). Human AAA blood were collected pre-surgery and tissue samples were collected and obtained from our local Biobank at the Department of Vascular- and Endovascular Surgery at University Hospital Düsseldorf (study number: 5731R and 2018-248-FmB).

### 2.3 Human platelet and plasma preparation

Platelet count and mean platelet volume (MPV) in blood from AMCs and AAA patients were analyzed using a hematology analyzer (Sysmex KX-21N, Norderstedt, Germany). For plasma and platelet preparation, blood was centrifuged at 231 *g* for 10 min. Afterwards, the upper phase, i.e. the platelet-rich plasma (PRP), was carefully transferred into Dulbecco’s phosphate buffered saline (DPBS, pH 6.5) containing apyrase (2.5 U/mL) (Sigma-Aldrich, #A7646) and Prostaglandin E1 (PGE_1_, 1 µM) (Sigma-Aldrich, #P5515). The DPBS PRP mixture was centrifuged at 1,000 *g* for 6 min without brakes. The formed pellet was resuspended in Tyrode’s buffer (137 mM NaCl, 2.8 mM KCl, 12 mM NaHCO_3_, 0.4 mM NaH_2_PO_4_ and 5.5 mM glucose, pH 6.5). Tyrode’s buffer is especially suitable to maintain the normal physiological function of isolated platelets. The cell count was determined using a hematology analyzer (Sysmex KX-21N, Norderstedt, Germany) and adjusted for the following experiment. For plasma sample preparation, the citrate tube was centrifuged again at 1,500 *g* for 10 min and at 4 °C after PRP extraction. The generated platelet-free plasma (PFP) was transferred into fresh collection tubes and stored at -70°C for later experiments.

### 2.4 Generation of platelet releasates

Isolated platelets (70 x 10^6^) from healthy volunteers were activated with ADP (10 µM, 10 min) and CRP (5 µg/mL, 5 min), control samples were treated with Tyrode’s buffer. The reaction was stopped by addition of apyrase (2 µL, 0.25 U/mL) and centrifuged at 850 *g* for 10 min. Finally, the activated platelet supernatant was collected and stored at -20 °C until use.

### 2.5 Experimental animals: The Pancreatic Porcine Elastase (PPE) mouse model

10-week-old standard C57BL/6J mice were purchased through Jackson or Janvier Laboratory. To induce experimental AAA, mice were anesthetized with 2-3% isoflurane and received a locally subcutaneous (s.c.) injection of buprenorphine (0.1 mg/kg) (Temgesic, Eumedica) 30 min before surgery. After median laparotomy, the proximal and distal infrarenal aorta were temporarily ligated to create an aortotomy above the iliac bifurcation. A catheter was inserted into the distal end of the aortotomy and used to infuse the aorta with sterile isotonic saline (NaCl, 0.9%) containing type I Porcine Pancreatic Elastase (2.5 U/mL) (Sigma-Aldrich, #E1250) at 120 mm Hg for 5 min. The vessels in control mice were infused with sodium chloride only. After removal of the infusion catheter, the aortotomy was closed without constriction of the aortal lumen. Finally, the abdomen was closed. For pain relief, all animals received Buprenorphine s.c. every 6 h during the daylight phase for three days and in the dark phase via drinking water (0.3 µg/mL). At the endpoints of the experiments (days 3, 10, 28) mice were euthanized under deep isoflurane anesthesia followed by cervical dislocation. For the platelet depletion experiments mice were injected with platelet depletion antibody (Emfret Analytics, polyclonal anti-GPIb alpha #R300) and a corresponding IgG antibody (Emfret Analytics, #C301) as controls after surgery (days 0 and 5). Endpoints of the experiments were days 3 and 10 post-(PPE)surgery.

### 2.6 Platelet preparation and total blood cell count in mice

Murine blood was collected and transferred into a heparin solution (20 U/mL). Platelet count and mean platelet volume (MPV) were determined using a hematology analyzer (Sysmex KX-21N, Norderstedt, Germany). For platelet preparation, the blood was centrifuged at 250 *g* at RT for 5 min. Then the supernatant was centrifuged again at 50 *g* for 6 min. To collect platelet-rich plasma (PRP), the supernatant of the last step was centrifuged with apyrase (0.02 U/mL) and prostacyclin (PGI_2_, 0.5 μM) at 650 *g* and RT for 5 min. Plasma samples were stored at -70°C for later experiments. The platelet pellet was resuspended and washed twice in murine Tyrode’s buffer (136 mM NaCl, 0.4 mM Na_2_HPO_4_, 2.7 mM KCl, 12 mM NaHCO_3_, 0.1% glucose, 0.35% BSA, pH 7.4) supplemented with PGI_2_ (0.5 μM) and apyrase (0.02 U/mL) by centrifugation at 650 *g* at RT for 5 min. Finally, platelets were resuspended in Tyrode’s buffer and incubated at 37 °C until use.

### 2.7 Transmission Electron Microscopy

Aortic tissue was harvested from PPE-operated mice. Briefly, the aorta was perfused with heparin (180 units/15 mL PBS) from the left ventricle (butterfly puncture) to right atrium (incision). Thereafter, perfusion was slowly carried out with 4% paraformaldehyde (in PBS pH 7.4). Afterwards, aortic tissue was fixed with a solution of 2% glutaraldehyde and 2% paraformaldehyde in cacodylate buffer (0.1 M, pH 7.2) at RT overnight. The next day, samples were washed twice in sodium cacodylate buffer (0.1 M) at RT for 10 min, post-fixed with 2% aqueous osmium tetroxide (OsO_4_) at RT for 5 h, washed twice with milliQ water and stained *en bloc* with 4% uranyl acetate overnight. After dehydration through a graded ethanol series and propylene oxide, the tissue was infiltrated and embedded in 100% eponate 12 resin (Ted Pella, Inc., Redding, CA, USA). Polymerization was performed at 60 °C for 24 h. Ultrathin sections were cut using a Leica Ultracut microtome (Leica, Deerfield, IL) and then post-stained with Reynolds lead citrate. The stained samples were examined in a FEI Tecnai G2 Spirit BioTWIN transmission electron microscope (Hillsboro, OR) operated at 80kV. Electron micrographs were taken with an AMT XR41 CCD camera from Advanced Microscopy Techniques, Corporation (Woburn, MA).

### 2.8 Flow cytometry

For flow cytometry of human platelets, whole blood (fresh citrate-anticoagulated blood) was used and diluted 1:10 in human Tyrode’s buffer (137 mM NaCl, 2.8 mM KCl, 12 mM NaHCO_3_, 0.4 mM NaH_2_PO_4_ and 5.5 mM glucose, pH 6.5). Heparinized mouse blood was washed twice at 250 *g* for 5 min by discarding the supernatant and resuspending the blood pellet in murine Tyrode’s buffer (136 mM NaCl, 0.4 mM Na_2_HPO_4_, 2.7 mM KCl, 12 mM NaHCO_3_, 0.1% glucose, 0.35 % BSA, pH 7.4). Finally, the pellet was diluted in mouse Tyrode’s buffer including CaCl_2_ (2 mM). Blood samples were mixed with antibodies and stimulated with indicated agonists at RT for 15 min in the dark. The reaction was stopped by the addition of DPBS and samples were analysed on a FACSCalibur flow cytometer (BD Biosciences).

Briefly, two-color analysis of murine or human platelet activation was performed using fluorophore-labeled antibodies for P-selectin expression (Emfret Analytics, Wug.E9-FITC, #M130-1, and BD Biosciences, anti-human CD62P-PE, #348107) and the active form of ⍺ _IIb_β_3_ integrin (Emfret Analytics, JON/A-PE, #M023-2, and BD Biosciences anti-human PAC-1-FITC, #340507). For analysis of α_5_ glycoprotein surface expression and up-regulation as well as activation of integrin β_3_ expression, blood samples were mixed with specific antibodies (Emfret Analytics, β_3_ chain, GPIIIa, Luc.H11-FITC, #M031-1 or Integrin ⍺ _5_/CD49e, Tap.A12-FITC and anti-human CD61-PE, #555754, BD Biosciences) and incubated for 15 min at RT in the dark. For the analysis of CD45- and Ly6G-platelet aggregates, specific antibodies were used (Emfret Analytics, GPIb/CD42b, Xia.G5-PE; BD Biosciences, anti-mouse CD45-APC, #559864 and BioLegend, anti-mouse Ly-6G-APC, #127614). For detection of phosphatidylserine (PS) exposure, Cy™5 Annexin V (BD Biosciences) were used – staining was performed while using binding buffer (10 mM Hepes, 140 mM NaCl, 2.5 mM CaCl_2_, pH 7.4) instead of DPBS. By treating platelets with soluble full-length OPN (0.01 µg/µL in H_2_O) (PeproTech, #120-35) and soluble cleaved OPN (0.01 µg/µL), the influence of osteopontin on platelet activation should be analyzed. Additionally, circulating levels of chemokines and other inflammatory mediators including CXCL8 (IL-8), CCL2 (MCP-1), CCL5 (RANTES), CXCL8 (IL-8), IL-6, P-selectin, PSGL-1, IL-1β, IFN-γ, IFN-α2, IL-10, IL-12p70 were determined in the plasma of AAA patients and age-matched controls (AMCs) by flow cytometric multiplex analysis (cytometric bead array-CBA) (Biolegend, LEGENDplex™ Human Thrombosis Panel (10-plex) #740892, and LEGENDplex™ Human Inflammation Panel 1 (13-plex) #740809 #740895) according to manufacturer’s instructions using a BD FACSLyric^TM^ flow cytometer.

### 2.9 Statistics

Data are shown as arithmetic means ± SEM, and statistical analysis was performed using GraphPad Prism Version 8.4.3. For each experiment sample size reflects the number of independent biological replicates and is provided in the figure legend. Statistical testing for mice was performed based on the assumption of normality because of low numbers of replicates and the use of inbred animals. Equality is part of the determination of statistical significances using GraphPad Prism 8 software and its recommended tests, which we applied throughout this study. Where necessary, clinical data were evaluated to check the normality of the distribution using the Shapiro-Wilk test.

Thus, data between two comparative groups, an unpaired student’s t-test with Welch’s correction or multiple t-test were used. For nonparametric data, a Mann-Whitney *U* test was used. Comparison of more than two groups was performed using a one- or two-way ANOVA with Sidak’s, Holm-Sidak’s or Tukey’s multiple-comparison correction. *P* values of less than 0.05 were considered significant.

## 3. Results

### 3.1 Platelet depletion alleviates inflammation and experimental AAA progression

We induced experimental murine AAA utilizing the porcine pancreatic elastase (PPE) model and performed platelet depletion by administrating an antiplatelet antibody (or control polyclonal immunoglobulins) at days 0 and 5 following surgery (**Figure 1A and B**). We found that platelet depletion significantly attenuated the diameter progression of AAA over a 10-day time course by ultrasound tracking (**Figure 1C-E, Supplemental Figure S1A**). Further, our data demonstrated enhanced aortic pulse wave velocity (PWV) following initiation of experimental AAA in the control group at day 10 post-surgery suggesting significant structural stiffening effects in the aortic wall. Of note, such changes in aortic PWV were not found in the platelet-depleted group, suggesting aggravating effects of platelets with adverse stiffness promoting structural remodeling (**Figure 1F**). In the thoracic aorta, stable media lamellar units and lack of elastic degradation were found in both groups (**see Supplementary Figure S1B-E**). In addition to the effects seen in the structural architecture of the AAA wall, our findings demonstrated that platelets and macrophages were recruited into the AAA wall at day 3 post-surgery. Platelet depletion reduced the number of macrophages in the AAA wall, suggesting potential platelet-mediated modulating effects of the inflammatory process following murine AAA induction (**Figure 1G-H, Supplemental Figure S1F and G**). In line with this finding, inflammatory gene expression analysis of whole AAA tissue explants revealed downregulation of major inflammatory genes in the platelet-depleted group versus the control antibody treated group at day 3 post-surgery (**Figure 1I**).

**Figure 1.**
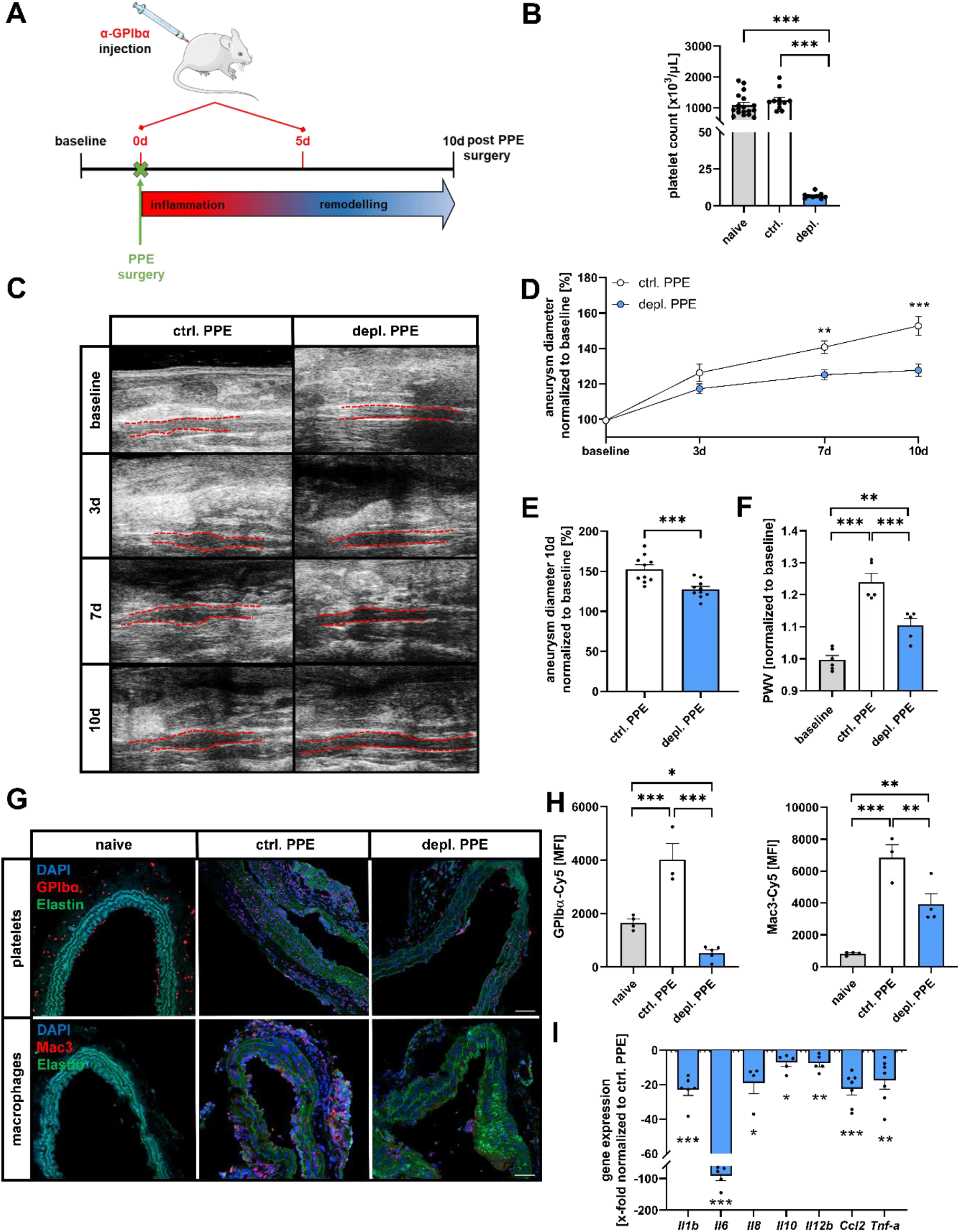
Platelet depletion reduces AAA progression and attenuates inflammatory response. (**A**) Schematic of experimental procedure: C57BL/6J mice received a platelet-depleting anti-GPIbα (depl.) or IgG control (ctrl.) antibody (at days 0 and 5) after PPE surgery. (**B**) Platelet counts of unoperated mice before and after platelet depletion (n = 11–18). (**C**) Ultrasound images at indicated time points (baseline, days 3, 7 and 10) of depleted and control mice after PPE surgery to investigate aneurysm development. (**D**) Aneurysm diameter was determined by ultrasound measurements at the indicated time points (n = 10). Data were normalized to baseline. (**E**) Aortic diameter progression of depl. PPE and ctrl. PPE mice at day 10 post-surgery, normalized to baseline (n = 10). (**F**) Calculated pulse wave velocity (PWV) 10 days after PPE surgery, normalized to baseline (n = 5–6). (**G**) Representative immunofluorescence (IF) images of platelets (CD42b/anti-GPIbα, red, upper panels), macrophages (anti-Mac3, red, lower panels), nuclei (DAPI, blue) and elastin (autofluorescence, green) in aortic tissue sections of depl. PPE and ctrl. PPE at day 3 post-surgery (n = 3–5). Scale bar: 50 µm. Aortic tissue of naive mice served as control (n = 4). (**H**) Quantification of platelet (GPIbα, left) and macrophage (Mac3, right) migration into the aortic tissue of depl. PPE and ctrl. PPE mice at day 3 post-surgery (n = 3–5). Naive mice served as control (n = 4). (**I**) Inflammatory gene expression in aortic tissue of depl. PPE mice 3 days after surgery. Fold changes were normalized to ctrl. PPE mice (n = 4–7). Data are represented as mean ± SEM. Statistical significances were determined by one-way ANOVA with Holm-Sidak’s multiple comparison (**B, F, H**), two-way ANOVA with Sidak’s multiple comparison test (**D**), unpaired student‘s t-test (**E**) or multiple t-test (**I**). \**P* < 0.05, \*\**P* < 0.01, \*\*\**P* < 0.001. Ccl, CC chemokine ligand; Ctrl, control; DAPI, 4′,6-diamidino-2-phenylindole; depl., depletion; Il, -interleukin; PPE, porcine pancreatic elastase (infusion); PWV, pulse wave velocity; Tnf, tumour necrosis factor.

### 3.2 Platelet depletion reverses adverse remodeling of the extracellular matrix

We next analyzed platelet-associated adverse remodeling effects on the extracellular matrix (ECM) of the AAA wall. Exemplary experiments show fragmented, and at given locations completely disrupted, elastin membranes within the tunica media in the AAA wall of PPE mice as shown by transmission electron microscopy (TEM) (**Figure 2A and B**). Therefore, we analyzed AAA wall fragmentation in more detail by hematoxylin/eosin and Verhoeff van Gieson staining. To examine the impact of platelets on elastin membrane integrity, histological evaluation of AAA explants was performed. Our results show that the intima media thickness of the AAA wall of platelet-depleted mice was thinner compared to control mice at 10 days post-surgery (**Figure 2C**). Additionally, the aortic tissue displayed significantly less fragmentation of elastin membranes in AAA explants of platelet-depleted versus control mice (**Figure 2D**).

**Figure 2.**
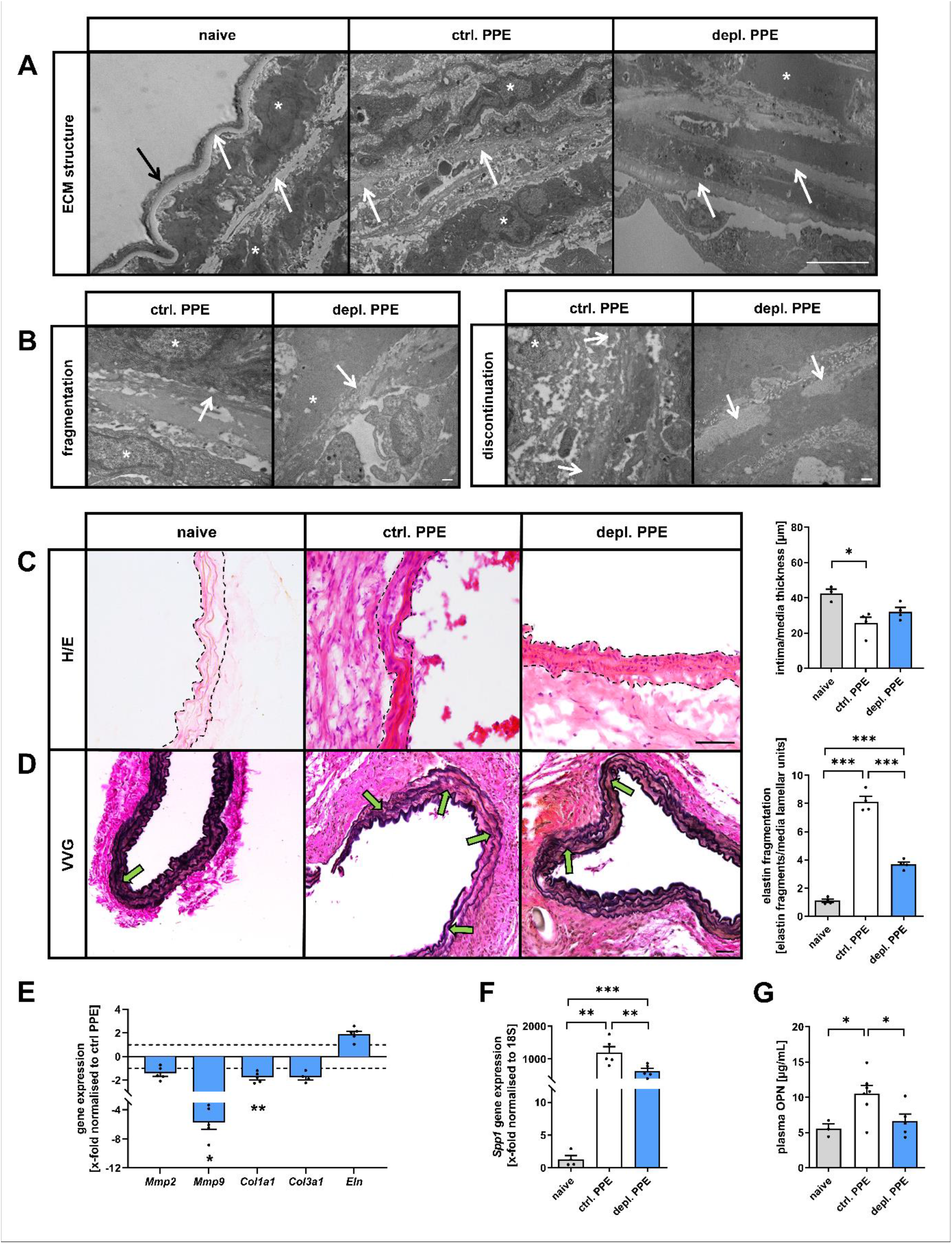
Platelet depletion alters ECM remodeling in PPE-operated mice 10 days after surgery. (**A**–**G**) C57BL/6J mice received a platelet depleting anti-GPIbα (depl.) or IgG control (ctrl.) antibody (at days 0 and 5) after PPE surgery and were sacrificed at day 10 for tissue preparation. (**A**) Exemplary Transmission electron microscopy (TEM) images of aortic tissue from depl. PPE and ctrl. PPE mice 10 days after surgery. Aortic tissue of naive mice served as control. (2,900x magnification, n = 2). Black arrow indicates elastin, white arrowheads indicate endothelial layer, and stars indicate vascular smooth muscle cells. Scale bars: 10 µm. (**B**) TEM images of fragmented and discontinuous aortic elastic tissue from depl. PPE and ctrl. PPE mice at day 10 post-surgery (13,000x magnification, n = 2). Scale bars: 0.5 µm. (**C** and **D**) Representative images of aortic tissue from naive and PPE-operated mice using (**C**) hematoxylin-eosin (H/E) staining and (**D**) Verhoeff van Gieson (VVG) staining. Intima/media thickness was determined after H/E staining (n = 3–4) and elastin fragmentation (breakdowns) was counted after VVG staining (n = 4). Scale bars: (**C**) 50 µm and (**D**) 100 µm. (**E**) Stiffness-related gene expression in aortic tissue of naive and PPE-operated mice. Fold change was normalized to ctrl. PPE mice (n = 4–5). (**G**) *Spp1* (OPN) gene expression in aortic tissue of naive and PPE-operated mice. Fold change was normalized to 18S housekeeping gene (n = 4–5). (**H**) Osteopontin plasma concentration (µg/mL) in naive and PPE-operated mice (n = 3–7). Data are represented as mean ± SEM. Statistical significances were determined by one-way ANOVA with Holm-Sidak’s multiple comparison test (**C**, **D**, **F** and **G**) and multiple t-test (**E**). \**P* < 0.05, \*\**P* < 0.01, \*\*\**P* < 0.001. Col, collagen; ctrl, control; depl., depletion; ECM, extracellular matrix; Eln, elastin; H/E, hematoxylin-eosin; Mmp, matrix metalloproteinase; OPN, osteopontin; PPE, porcine pancreatic elastase (infusion); *Spp1*, secreted phosphoprotein 1; VVG, Verhoeff van Gieson.

These morphologic findings were supported by a partially altered structural gene expression profile in AAA of platelet-depleted mice. In detail, AAA whole tissue explants of platelet-depleted mice showed reduced expression levels of matrix metalloproteinase (*Mmp*) *9* and *collagen I*, while *collagen III*, *Mmp2* and *elastin* (*Eln*) expression were unaltered (**Figure 2E**). MMP9 is considered to be a major elastolytic enzyme that reduces resilience and elasticity of conduit arteries and was downregulated in whole AAA explants of platelet-depleted versus control mice 10 days post-surgery (**Figure 2E**), consistent with the aforementioned reduced elastin fragmentation in the same group. Notably, we found that osteopontin (*Spp1*/OPN), a multifunctional and highly phosphorylated glycoprotein was up-regulated at the gene expression level in aortic tissue of PPE-operated mice and in the plasma following murine experimental AAA induction. *Spp1* expression was down-regulated within the AAA wall of platelet-depleted mice compared to control mice, and OPN protein was reduced in the plasma (**Figure 2F and G**).

### 3.3 Platelets alter gene expression in macrophages

Since OPN was altered at the transcriptional and protein level upon platelet-depletion, we elected to study the major source of OPN. We incubated macrophages (PMA-stimulated THP-1 cells; differentiated macrophages), aortic vascular smooth muscle cells (hASMC) and aortic fibroblasts (hAoF) with the releasates of ADP (stimulating P2Y_12_) or CRP (collagen related peptide, stimulating the collagen receptor GPVI) activated platelets. Our results demonstrated that macrophages are the major cell type with the highest *SPP1* expression when compared to resident cells of the AAA wall, such as hASMC and hAoF (**Figure 3A**). Although platelets themselves carry OPN (**Figure 3B**), differentiated macrophages remained the primary source for OPN protein and *SPP1* gene expression *in vitro* (**Figure 3C and D, Supplemental Figure S2A**). To verify whether platelets are capable of altering gene expression in macrophages, we exposed cultures of pro-inflammatory macrophages to releasates of activated platelets (with ADP and CRP) *in vitro* (**Figure 3E and F**). The data revealed that platelet releasates alter inflammatory gene expression in macrophages. Activated platelet releasates caused marked up-regulation of key inflammatory genes, such as Interleukins *(IL)-6*, -*12B* and -*1B* but only moderate gene expression of TNF-α or IL-8 (**Figure 3E**). We next studied the influence of releasates on stiffness-related gene expression in macrophages. Releasates from activated platelets enhanced gene expression of *SPP1, MMP9 and collagen Ia1* in macrophages (**Figure 3F**), which supports our *in vivo* findings and suggests that a crosstalk between platelets and macrophages may be a central mechanism for AAA initiation and diameter progression. In addition, we could show for the first time that treatment of hAoFs with releasates of CRP-activated platelets induced IL-6 gene expression and IL-6 release from these cells (**Figure 3G and H**). These results strongly suggest a significant contribution of platelets in paracrine effects of fibroblasts that amplify monocyte recruitment and activation in the pathogenesis of AAA^20^. Next, we also analyzed platelet-induced MMP9 expression of hAoFs by incubating the cells with the supernatant of ADP and CRP activated platelets. As shown in **Figure 3I**, we detected MMP9 expression also in hAoFs when incubated with the supernatant of activated platelets. However, according to the current literature, MMP9 gene expression was low in AoFs compared to macrophages (**Figure 3I**)^21^.

**Figure 3.**
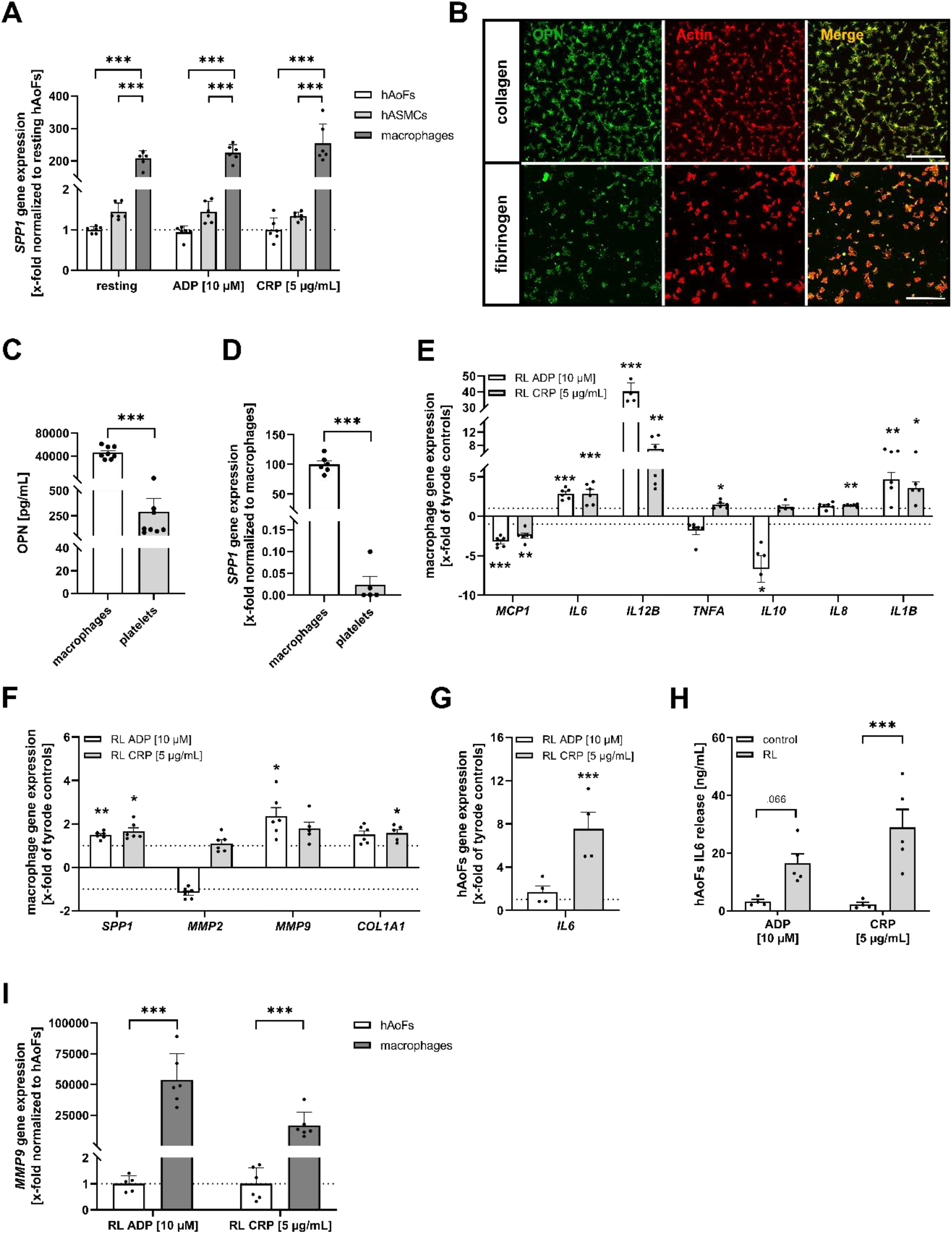
Platelet releasates affect inflammatory and stiffness-related gene expression of macrophages *in vitro.* (**A**) *SPP1* gene expression of hAoFs, hASMCs and differentiated macrophages (derived from PMA activated THP-1 cells) in cell culture experiments stimulated with releasates (RL) of human platelets for 6 h (n = 6). Platelets were stimulated with indicated agonists. Fold changes of hASMCs and differentiated macrophages were normalized to hAoFs treated with resting platelet releasates. (**B**) Representative fluorescence images of OPN (anti-OPN, green) in adherent platelets (actin staining with rhodamine-phalloidin, red) on collagen (200 µg/mL) and fibrinogen (100 µg/mL) after 20 min (n = 3). Scale bar: 50 µm. (**C**) OPN release of differentiated macrophages and ADP (10 µM) stimulated platelets was determined by ELISA (n = 8). (**D**) *SPP1* gene expression in differentiated macrophages and resting platelets (n = 5–6). (**E** and **F**) Inflammatory and stiffness-related gene expression of differentiated macrophages, incubated with releasates of human platelets (RL) for 6 h. Platelets were stimulated with indicated agonists. Fold change was normalized to corresponding tyrode controls (n = 4–6). (**G**) *IL6* gene expression of hAoFs, incubated with releasates of human platelets (RL) for 24 h. Platelets were stimulated with indicated agonists. (n = 4). (**H**) IL6 release of hAoFs, incubated with releasates of human platelets (RL) for 24 h. Platelets were stimulated with indicated agonists (n = 4–5). (**I**) *MMP9* gene expression of hAoFs and macrophages in cell culture experiments stimulated with releasates (RL) of human platelets for 6 h (n = 5–6). Platelets were stimulated with indicated agonists. Fold changes were normalized to hAoFs gene expression. Data are represented as mean ± SEM. Statistical significances were determined by two-way ANOVA with Sidak’s multiple comparison (**A, H, I**), unpaired student‘s t-test (**C, D** and **G**) or multiple t-test (**E** and **F**). \**P* < 0.05, \*\**P* < 0.01, \*\*\**P* < 0.001. ADP, adenosine diphosphate; Col, collagen; CRP, collagen-related peptide; hAoFs, human aortic fibroblasts; hASMCs, human aortic smooth muscle cells; MCP, monocyte chemoattractant protein; MMP, matrix metalloproteinase; OPN, osteopontin; Il, interleukin; RL, releasate; *SPP1*, secreted phosphoprotein 1; TNF, tumour necrosis factor.

### 3.4 Platelets adhere to immobilized osteopontin via integrin α_V_β_3_ under static and dynamic conditions

To determine the impact of OPN in platelet-mediated responses in inflammation and aortic wall remodeling in AAA, we analyzed the expression of OPN in aortic explants. While only limited OPN expression was detected in the aortic wall of naive mice, we found elevated OPN protein expression in AAA explants 28 days post-surgery (**Figure 4A**). To analyze a potential role for aortic OPN in recruiting platelets to the vessel wall in AAA, we determined platelet adhesion to immobilized full-length and cleaved OPN under static conditions. Cleavage of OPN was achieved by thrombin treatment and verified by silver staining (**see Supplementary Figure S2B**). Activation of human platelets with ADP and CRP resulted in enhanced platelet adhesion to full-length and cleaved OPN with a similar amplitude (**Figure 4B**). Control experiments examined platelet adhesion to fibrinogen (positive control) and BSA (negative control), and revealed that platelet adhesion on OPN is highly up-regulated after platelet activation. Incubation of platelets with the α_V_β_3_ inhibiting antibody MAB1976 revealed that platelets adhere to OPN – at least partially– through integrin α_V_β_3_ because platelet adhesion to OPN was significantly reduced compared to controls (**Figure 4C**).

**Figure 4.**
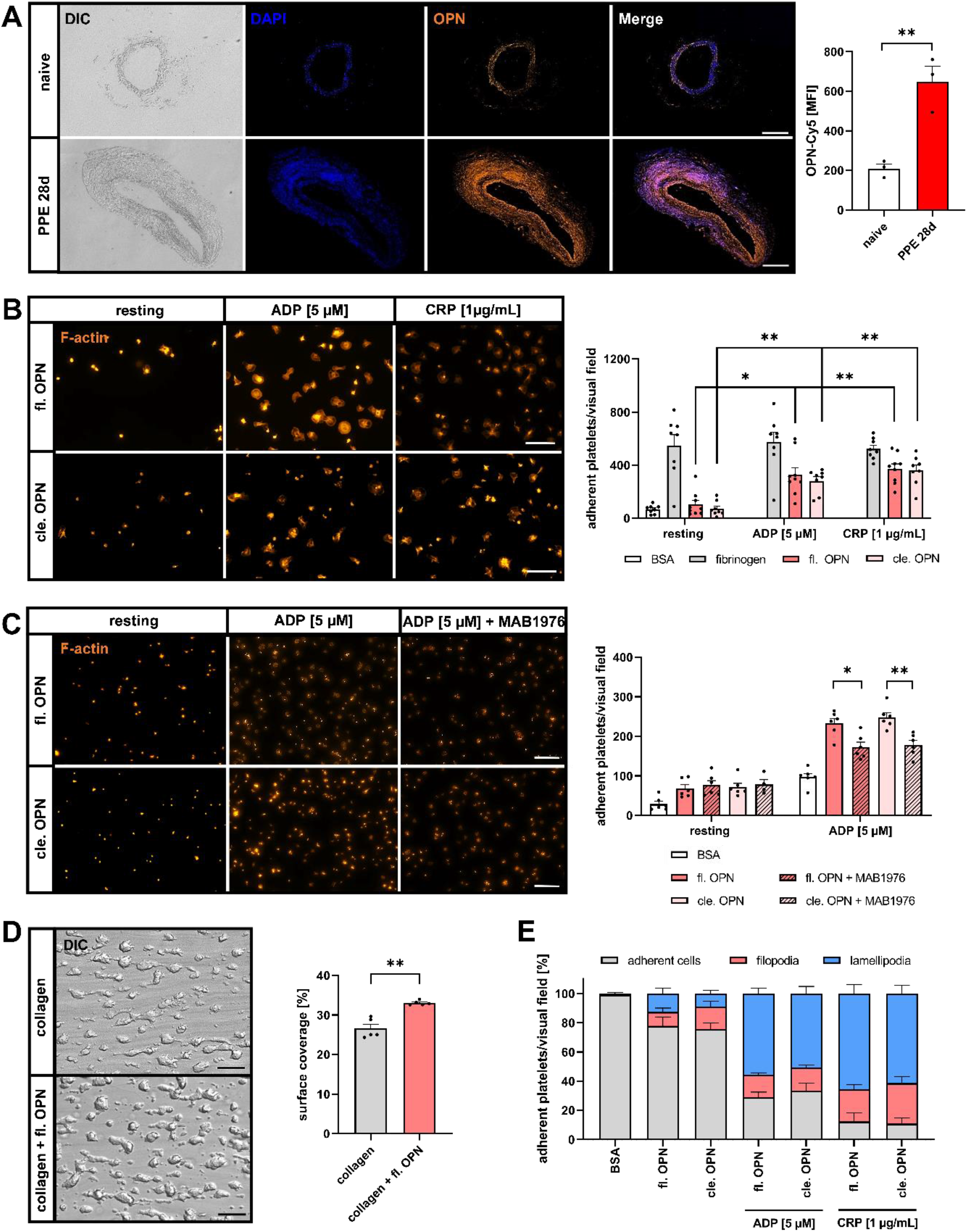
Platelets adhere to immobilized full-length osteopontin via integrin α_V_β_3_ under static and dynamic conditions. (**A**) Representative immunofluorescence (IF) images and quantification of aortic tissue from naive and PPE-operated mice at day 28 post-surgery, specifically stained for osteopontin (anti-OPN, orange) and nuclei (DAPI, blue) (n = 3). Scale bar: 50 µm. (**B**) Representative images and quantification of adherent human platelets stimulated with indicated agonists, on BSA (1%), fibrinogen (100 µg/mL), cleaved or full-length osteopontin (5 µg/mL) after 30 minutes. Adherent platelets were visualized by rhodamine-phalloidin staining (F-actin, orange) (n = 5). Scale bar: 20 µm. (**C**) Representative images and quantification of adherent resting and activated (ADP 10 µM) human platelets, on BSA (1%), cleaved or full-length osteopontin (5 µg/mL) after treatment with or without MAB1976 (α_V_β_3_ inhibiting antibody) for 30 min (n = 5). Scale bar: 50 µm. (**D**) Representative images and quantification of thrombus formation experiments with human whole blood on a collagen matrix (200 µg/mL) and a collagen/full-length osteopontin matrix (collagen: 200 µg/mL, fl. OPN: 100 µg/mL) at an arterial shear rate of 1700 s_-1_ (n = 5). Scale bar: 50 µm. (**E**) Number of filopodia- and lamellipodia-forming platelets spread on different matrices, indicated as percent of adherent cells per visual field (n = 5). Data are represented as mean ± SEM. Statistical significances were determined by two-way ANOVA with Tukey’s multiple comparison (**B** and **C**) or unpaired student‘s t-test (**D**). \**P* < 0.05, \*\**P* < 0.01. ADP, adenosine diphosphate; BSA, bovine serum albumin; cle., cleaved; CRP, collagen-related peptide; DAPI, 4′, 6-diamidino-2-phenylindole; DIC, differential interference contrast; fl., full-length; OPN, osteopontin; PPE, porcine pancreatic elastase (infusion).

Under high arterial shear conditions, we next examined thrombus formation on a collagen matrix and compared the surface coverage of three-dimensional thrombi with platelet adhesion and aggregate formation on a mixed collagen-OPN matrix using a well-established *ex vivo* flow chamber system. As shown in figure 4D, increased thrombus formation was observed on the collagen-OPN matrix compared to collagen alone, suggesting that OPN not only affects platelet adhesion but also thrombus formation under high arterial shear flow conditions found in the aorta (**Figure 4D**). Platelet adhesion to immobilized OPN induced the formation of filopodia and lamellipodia in platelets indicating that platelet adhesion to OPN affects cytoskeletal remodeling and increases the platelet surface area. Again, no major alterations in cytoskeletal remodeling were detected between platelets adhering to full-length and cleaved OPN. However, platelet activation by ADP or CRP strongly supported the formation of lamellipodia of platelets on the OPN matrix (**Figure 4E, see Supplementary Figure S2C**). When we analyzed platelet adhesion to collagen-full length OPN or collagen-cleaved OPN surfaces and compared them to collagen alone, we did not detect any significant changes under static conditions (**see Supplementary Figure S2D**). Furthermore, platelet adhesion was unaltered when platelets were pre-treated with soluble full-length or cleaved OPN (**see Supplementary Figure S2E**), suggesting that arterial shear conditions are crucial for amplified platelet adhesion and thrombus formation on a collagen-OPN matrix. To examine if soluble OPN is able to amplify platelet activation, we determined active integrin α_IIb_β_3_ and P-selectin exposure by flow cytometry. However, our results revealed no impact of soluble OPN on platelet activation following treatment with standard agonists such as ADP and CRP (**see Supplementary Figure S2F and G**).

### 3.5 Platelet activation and pro-coagulant activity are key elements for the progression of experimental AAA

Next, we examined if aberrant platelet activation is responsible for the observed platelet-mediated effects on inflammation and remodeling in AAA. To this end, we determined platelet integrin α_IIb_β_3_ activation and P-selectin exposure as marker for degranulation using flow cytometry. As shown in figure 5A, blood was drawn at various time points (days 3, 10 and 28 post-surgery) (schematic overview, **Figure 5A**) and aneurysm formation was monitored over a time period of 28 days after PPE surgery and compared to sham-operated controls (**Figure 5B, Supplemental Figure S3A**). At an early time point (day 3), slightly reduced P-selectin exposure and integrin activation but unaltered phosphatidylserine (PS) exposure were observed following platelet stimulation with PAR4 peptide that activated the thrombin receptor PAR4 (**Figure 5C**). At day 10 post PPE surgery, no differences were detected in the activation or pro-coagulant activity in platelets from PPE-operated mice compared to sham-operated controls (**Figure 5D**). However, at a later time point (day 28), we detected significantly enhanced P-selectin exposure and integrin α_IIb_β_3_ activation after stimulation of the collagen receptor GPVI using CRP. In addition, pro-coagulant activity of platelets isolated from PPE-operated mice was enhanced compared to sham-operated controls (**Figure 5E**).

**Figure 5.**
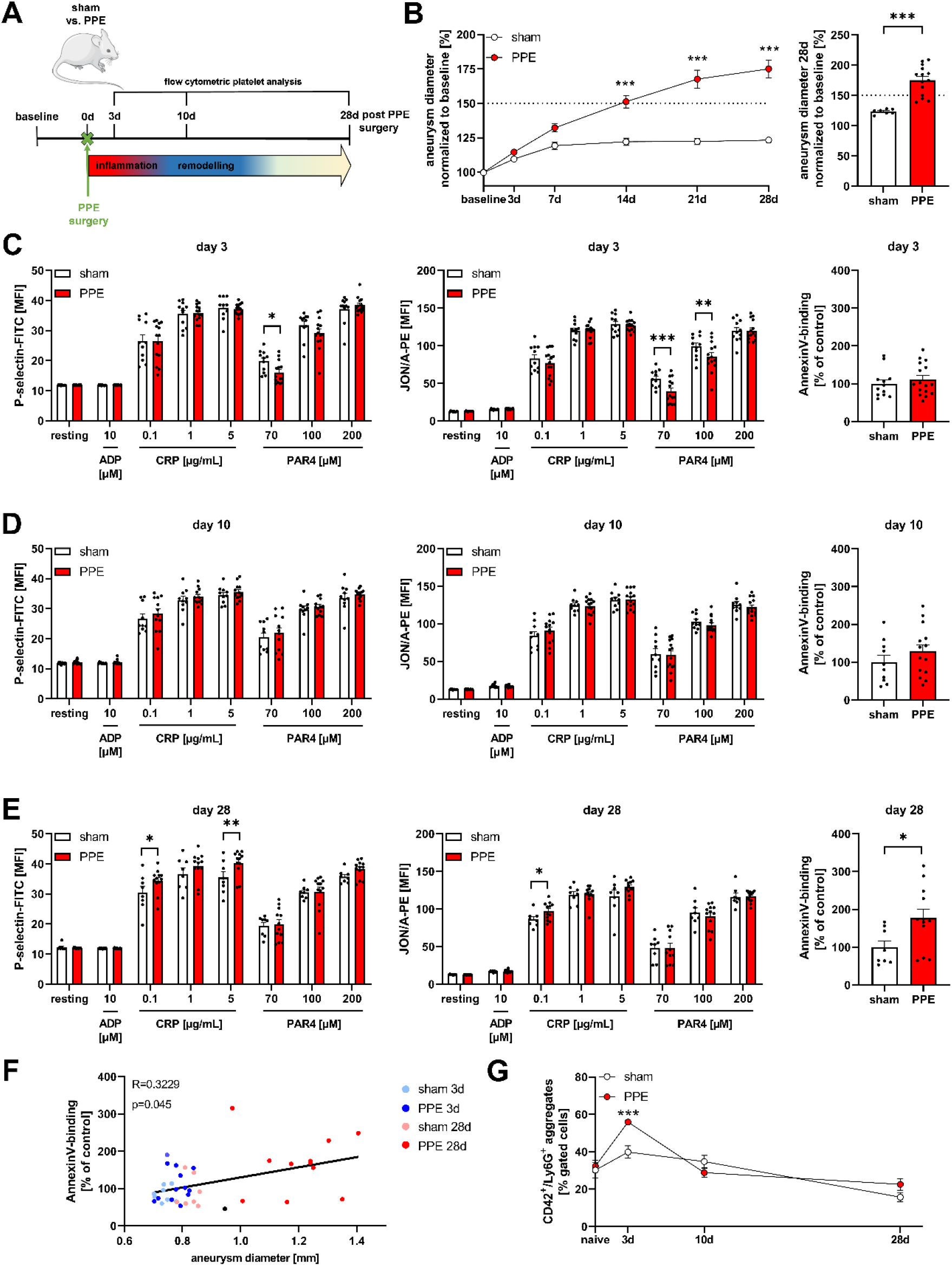
Platelet activation and pro-coagulant activity is crucial for the AAA progression in the murine PPE model. (**A**) Schematic of experimental procedure: C57BL/6J mice underwent sham or PPE surgery. Mice were sacrificed at indicated time points for flow cytometric analysis of platelets in whole blood at days 3, 10 and 28. (**B**) Ultrasound measurements were performed prior surgery (baseline) and at days 3, 7, 14, 21 and 28 to investigate aneurysm diameter (n = 8–13). Data were normalized to baseline (**C**–**E**) Platelet degranulation (P-selectin-FITC), active integrin α_IIb_β_3_ externalisation (JON/A-PE) and AnnexinV-binding to platelets from sham or PPE-operated mice at days 3 (**C**), 10 (**D**) and 28 (**E**) after PPE surgery were determined by flow cytometry. Platelets were stimulated with indicated agonists (n = 7–15). Data are represented as MFI or percent-gated cells normalized to control. (**F**) Spearman’s correlation between the diameter of the aneurysm and AnnexinV-binding at days 3 and 28 after PPE surgery (n = 8–15). (**G**) Aggregate formation of platelets (CD42_+_) and neutrophils (Ly6G_+_) in whole blood of naive, sham or PPE-operated mice at days 3, 10 and 28 after surgery determined by flow cytometry (n = 5–6). Data are represented as mean ± SEM. Statistical significances were determined by two-way ANOVA with Sidak’s multiple comparison test (**B** and **G**), unpaired student‘s t-test or multiple t-test (**C**, **D** and **E**). \**P* < 0.05, \*\**P* < 0.01, \*\*\**P* < 0.001. ADP, adenosine diphosphate; CRP, collagen-related peptide; MFI, mean fluorescence intensity; PAR4, protease-activated receptor 4 activating peptide; PPE, porcine pancreatic elastase (infusion).

Determination of Spearman’s correlation coefficient between the diameter of the aneurysm and Annexin V binding to platelets at days 3 and 28 after PPE surgery revealed a strong correlation between aortic diameter expansion and elevated pro-coagulant activity (**Figure 5F**) emphasizing a role for PS at the platelet surface in AAA pathology. It is worth mentioning that the number and size of platelets (**see Supplementary Figure S3B and C**), and the exposure of integrins on the platelet surface under resting and stimulated conditions (**see Supplementary Figure S3D and E**) were not altered at days 3, 10 and 28 after induction of AAA when compared to sham-operated controls. Furthermore, elevated platelet-neutrophil conjugates were detected in PPE-operated mice at day 3 post-surgery compared to sham-operated controls (**Figure 5G**) while no alterations were determined when we analyzed leukocyte-platelet conjugates (**see Supplementary Figure S3F**).

### 3.6 Platelet depletion reduces aortic diameter progression in experimental angiotensin-II induced AAA formation

To confirm the importance of platelets in AAA formation and progression in another experimental mouse model of AAA, we depleted platelets in Ang-II infused apolipoprotein E (ApoE) knock-out mice. We found that platelet depletion significantly attenuated the diameter progression of AAA over a 7-day time course by ultrasound tracking (**Figure 6A-B, see Supplemental Figure 4C**). In line with platelet depletion in PPE mice **(Figure 2C-D)**, we detected unaltered intima/media thickness but reduced elastin fragmentation in platelet depleted Ang-II infused ApoE mice **(see Supplemental Figure S4A-B)**. Next, we examined if this is due to aberrant platelet activation as observed in PPE mice. First, we confirmed aneurysm formation and progression in Ang-II infused ApoE mice compared to sham controls **(Figure 6C, see Supplemental Figure S4D)**. In Ang-II infused ApoE knock-out mice, significantly reduced P-selectin exposure and integrin α_IIb_β_3_ activation in response to different concentrations of CRP and PAR4 peptide were observed at an early time point (day 3) (**Figure 6D**). Furthermore, agonist induced up-regulation of integrin α_IIb_β_3_ at the platelet surface was reduced as well. In contrast to PPE-induced AAA formation, no major alterations of platelet activation were detected in Ang-II infused ApoE knock-out mice at later time points (day 10 and 28) (**Figure 6E-F, see Supplemental Figure S4E**). Pro-coagulant activity of platelets from Ang-II infused ApoE knock-out mice was unaltered at all-time points examined **(Figure 6E-F)**. In control experiments, we confirmed unaltered platelet counts and size in ApoE knock-out mice **(see Supplemental Figure S4F-G**).

**Figure 6.**
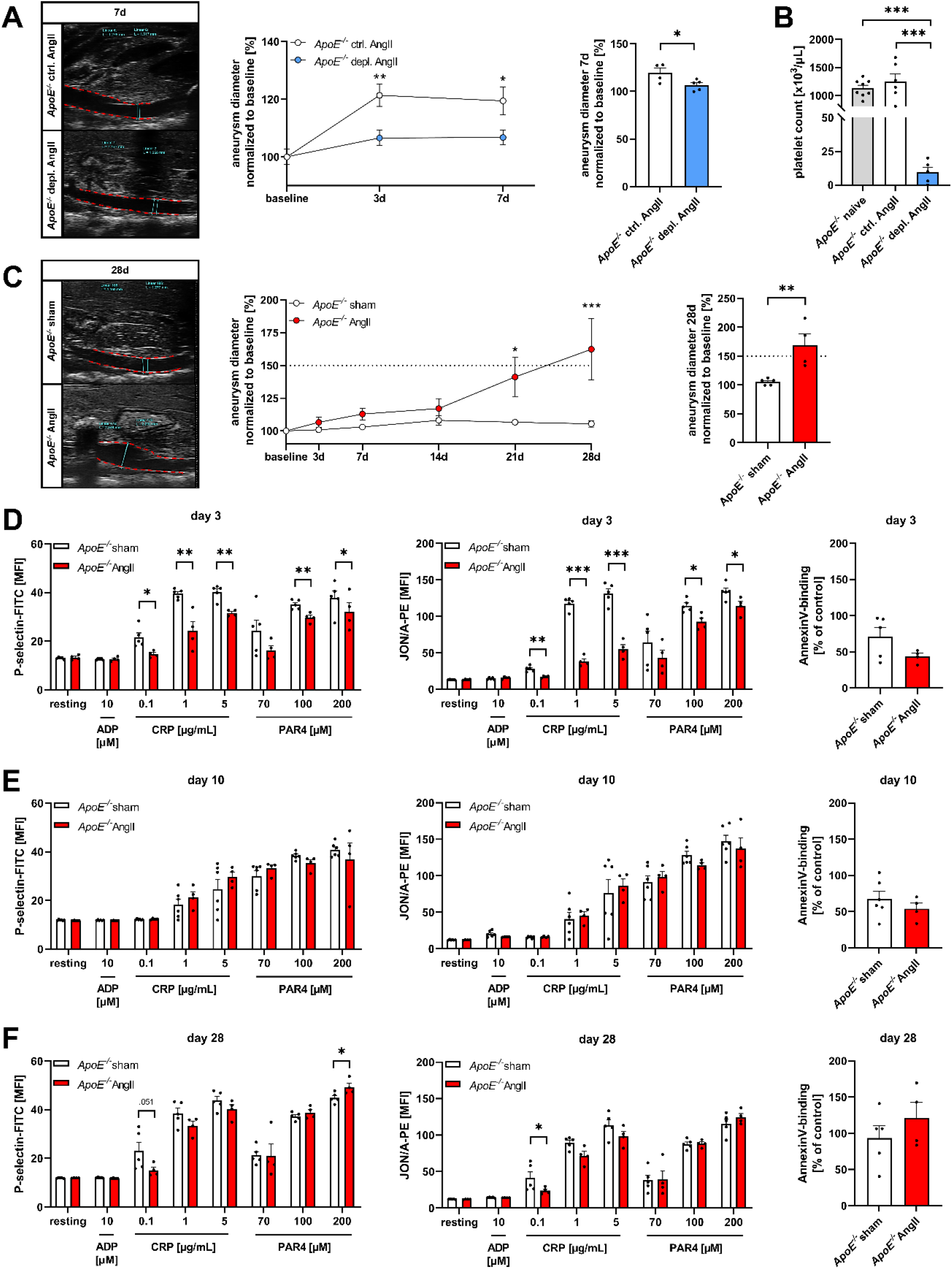
Platelet depletion reduces aortic diameter progression in experimental angiotensin-II induced AAA formation. (**A**) *ApoE_-/-_* mice received a platelet depleting anti-GPIbα (depl.) or IgG control (ctrl.) antibody (at days 0 and 5) after osmotic mini-pump implantation. Aortic diameter progression of depl. *ApoE_-/-_* and ctrl. *ApoE_-/-_* mice infused with Ang-II was measured via ultrasound over a time-period of 7 days (n = 4–5). Data were normalized to baseline. (**B**) Platelet counts of naive, depl. and ctrl. *ApoE_-/-_* mice before and after platelet depletion (n = 4–6). (**C**) Aortic diameter progression in sham or Ang-II infused *ApoE_-/-_* mice over a time-period of 28 days was determined via ultrasound measurements (n = 4–5). Data were normalized to baseline. (**D**–**F**) Platelet degranulation (P-selectin-FITC), active integrin α_IIb_β_3_ externalisation (JON/A-PE) and AnnexinV-binding to platelets in sham or Ang-II infused *ApoE_-/-_* mice at 3 (**C**), 10 (**D**) and 28 days (**E**) after osmotic mini-pump implantation were analysed via flow cytometry. Platelets were stimulated with indicated agonists (n = 4–5). Data are represented as MFI or percent-gated cells normalized to control. Statistical significances were determined by two-way ANOVA with Sidak’s multiple comparison test (**A** and **C**), unpaired student‘s t-test (**A** and **C**), one-way ANOVA with Holm-Sidak’s multiple comparison test (**B**) or multiple t-test (**D**, **E** and **F**). \**P* < 0.05, \*\**P* < 0.01, \*\*\**P* < 0.001. AAA, abdominal aortic aneurysm; ADP, adenosine diphosphate; Ang-II, angiotensin-II; CRP, collagen-related peptide; ctrl, control; depl., depletion; MFI, mean fluorescence intensity; PAR4, protease-activated receptor 4 activating peptide.

### 3.7 Platelet activation, pro-coagulant activity and OPN protein expression are important pathological features in patients with AAA

To study the relevance of platelet activation on inflammation in patients with AAA, we first determined integrin α_IIb_β_3_ activation and P-selectin exposure of platelets from AAA patients. As shown in figure 7A, we found elevated platelet activation in resting platelets, suggesting a pre-activated state of platelets in the blood circulation of AAA patients. Stimulation of the platelet collagen receptor GPVI reveals that the threshold where CRP is able to activate platelets is reduced in patients with AAA compared to age-matched healthy controls, as elevated platelet activation was detectable even with low-dose CRP (0.1 µg/mL). In addition, we detected elevated integrin α_IIb_β_3_ activation after ADP stimulation of platelets (**Figure 7A, see Supplemental Figure S5A**). Furthermore, reduced platelet counts but unaltered platelet size (MPV, mean platelet volume) were detected in AAA patients compared to age-matched controls (**see Supplementary Figure S5B**). Clinical characteristics of the AAA patient cohort are shown in Supplemental table 1.

**Figure 7.**
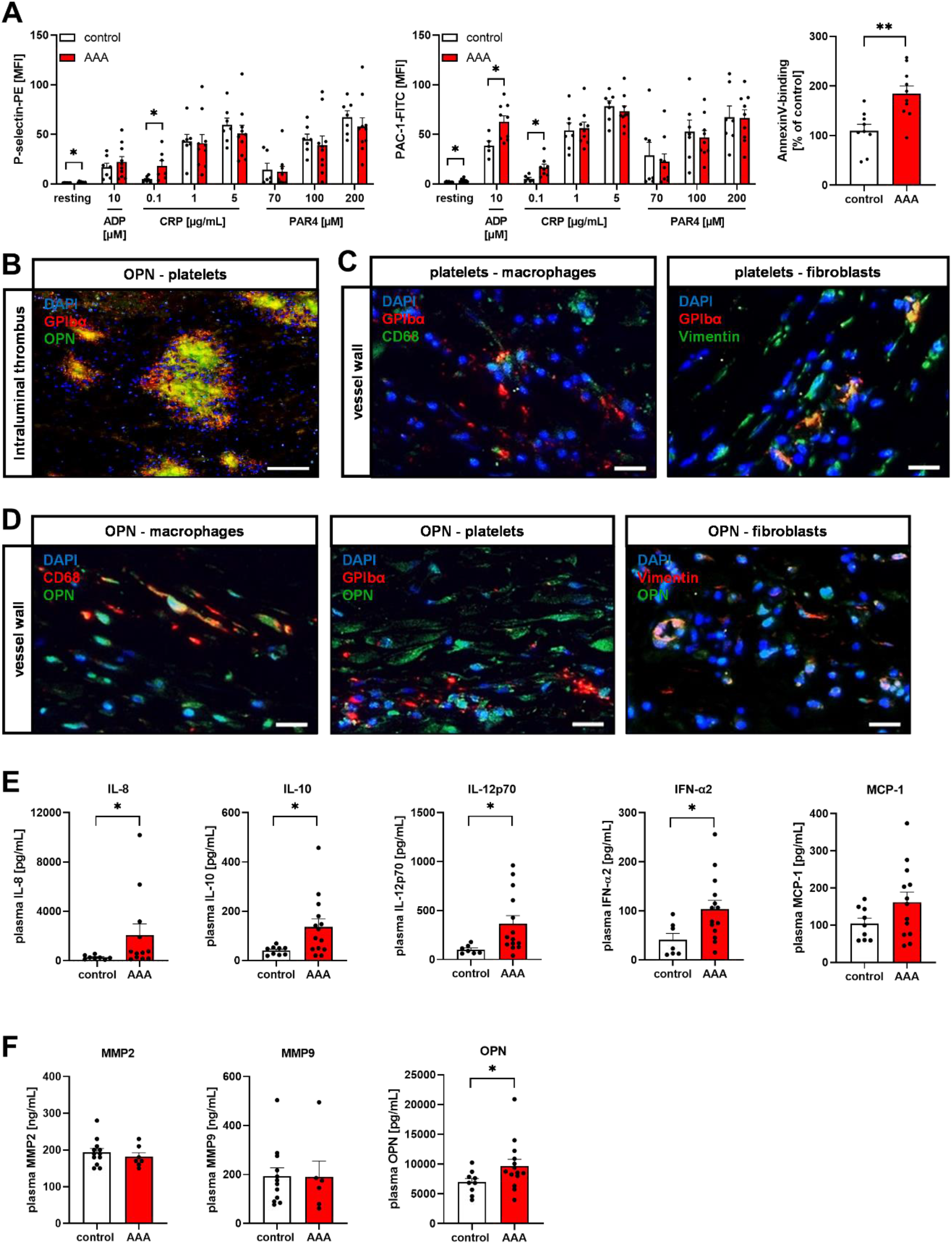
Platelet activation, pro-coagulant activity and osteopontin are important in AAA pathology. (**A**) Platelet degranulation (P-selectin-PE), active integrin α_IIb_β_3_ externalisation (PAC-1-FITC) and AnnexinV-binding to platelets in whole blood of AAA patients and age-matched controls were determined by flow cytometry (n = 5–10). Data are represented as MFI or percent-gated cells normalized to control. (**B**) Representative immunofluorescence images of platelets (anti-CD42b/GPIbα, 2 µg/mL, red), osteopontin (anti-OPN, 20 µg/mL, green) and nuclei (DAPI, blue) in the luminal layer of intraluminal thrombi (n = 5) of AAA patients. Scale bar: 100 µm. (**C**) Immunofluorescence staining of platelets (anti-CD42b/GPIbα, 2 µg/mL, red), nuclei (DAPI, blue) and macrophages (anti-CD68 AA 21-120, 10 µg/mL, green) respectively fibroblasts (anti-vimentin, 20 µg/mL, green) in aortic vessel wall of AAA patients. Scale bar: 20 µm. (**D**) Doublestaining of osteopontin (anti-OPN, 20 µg/mL, green) and nuclei (DAPI, blue) with macrophages (anti-CD68, 10 µg/mL, red), platelets (anti-CD42b/GPIbα, 2 µg/mL, red) and fibroblasts (anti-vimentin, 20 µg/mL, red) in aortic vessel wall. Scale bar: 20 µm. (**E**) Plasma concentration of circulating chemokines and inflammatory markers of AAA patients and age-matched controls were determined by flow cytometric multiplex analysis (cytometric bead array) (n = 7–14). (**F**) Plasma concentration of MMP2, MMP9 and OPN in AAA patients and age-matched controls using ELISA (n = 7–14). Data are represented as mean ± SEM. Statistical significances were determined by multiple t-test and unpaired student’s t-test (**A**) or unpaired t-test with Welch’s correction (**E** and **F**). \**P* < 0.05, \*\**P* < 0.01. AAA, abdominal aortic aneurysm; ADP, adenosine diphosphate; CRP, collagen-related peptide; DAPI, 4′, 6-diamidino-2-phenylindole; IFN, interferon; IL, interleukin; MCP, monocyte chemoattractant protein; MMP, matrix metalloproteinase; OPN, osteopontin; PAR4, protease-activated receptor 4 activating peptide.

Next, we analyzed aortic tissue and the ILT of AAA patients who underwent open surgical repair. Analysis of the ILT of different patients revealed that platelets and OPN protein accumulate and – more importantly – co-localize in the ILT suggesting that OPN protein accumulation might also modulate platelet recruitment, platelet adhesion and accumulation in the ILT (**Figure 7B and see Supplementary Figure S6B-C**). In line with the results from experimental AAA, we identified platelets in the aortic wall of patient specimens, suggesting platelet migration into AAA tissue (**Figure 7C-D, see Supplementary Figure S6A**). Furthermore, platelets co-localize to macrophages and fibroblasts in the vessel wall suggesting a crosstalk of these cells upon AAA progression in patients (**Figure 7C**). Next, we detected high OPN protein expression in the abdominal aortic wall of patient specimens, suggesting that OPN might be -at least in part-responsible for the recruitment of platelets to aortic tissue (**Figure 7D see Supplementary Figure S6A**). This is strengthened by the observed co-localization of platelets and OPN (**Figure 7D**).

Furthermore, we detected co-localization of macrophages and fibroblasts with OPN suggesting that these cells might be recruited to the aortic wall by OPN as well **(Figure 7D).** In addition to enhanced platelet activation and platelet recruitment to the aortic wall and the ILT, we detected elevated levels of inflammatory and anti-inflammatory cytokines such as IL-8, IL-10, IL-12 and IFN-α in the patient plasma (**Figure 7E**). In contrast, no alterations were detected in the plasma of AAA patients with regard to soluble P-selectin, PSGL-1, RANTES, IL-1β, IL-6 and INF-γ (**see Supplementary Figure S5C and D**). Furthermore, plasma levels of MMP2 and MMP9 were unaltered in AAA patients (**Figure 7F**). However, we again detected elevated OPN plasma levels in these patients, suggesting an important role of OPN for inflammation and remodeling of the abdominal aortic wall (**Figure 7F**).

## 4. Discussion

In this study, we found that platelets play a crucial role in AAA, likely due to the modulation of inflammation and degradation of the ECM induced by the functional crosstalk with macrophages and fibroblasts. Platelet depletion in experimental AAA reduced inflammation and aortic remodeling, leading to significantly reduced aortic diameter expansion in these mice. In AAA patients and in experimental mice, platelet activation and pro-coagulant activity was enhanced and might be responsible for platelet-induced effects on AAA pathology. Furthermore, we detected platelets and OPN in the vessel wall and in the ILT of patients who underwent open repair of AAA. Of note, OPN co-localized with platelets, suggesting a potential role for OPN in the recruitment of platelets to the ILT and probably to the aortic wall.

AAA is a multifactorial disease, which is characterized by chronic inflammation and degradation of the ECM in the abdominal aortic wall, often accompanied by the formation of an ILT. Increased platelet activation and aggregation is a characteristic feature of many cardiovascular diseases (CVDs), increasing the risk for thromboembolic events^13,22–25^. Here, we provide evidence for enhanced platelet activation at later time points in mice with experimental AAA (PPE-operated mice) and in patients with AAA. Both, PPE-operated mice and AAA patients exhibit elevated platelet responses after stimulation of the major collagen receptor GPVI using CRP, especially with low concentrations of the agonist. This, together with elevated platelet activation in resting, non-stimulated circulating platelets suggests that the threshold to induce platelet responses is reduced in experimental mice and in patients with AAA. The chronic inflammatory state in AAA might be a decisive determinant in priming circulating platelets for subsequent pro-coagulant and pro-thrombotic activity.

Recently, enhanced platelet reactivity (only P-selectin, no integrin activation) has been detected following thrombin and thromboxane receptor activation using platelets from AAA patients and experimental mice^26^. In contrast to our study, the authors used human platelets from healthy participants at any age as controls (not age-matched, although platelets behave differently with increasing age) and murine platelets isolated from a mouse model with topical application of elastase to induce AAA. In addition, they analyzed the thrombin/thromboxane receptor pathway in murine and human platelets but did not stimulate platelets with ADP or collagen/CRP to activate the major collagen receptor GPVI. Another study from Robless and colleagues detected elevated spontaneous platelet aggregation in AAA patients and in patients with peripheral artery disease (PAD), suggesting enhanced platelet activation even in the absence of agonists^27^. In contrast with our data, no differences were observed in PAC-1 binding or P-selectin exposure in resting platelets of patients with AAA or PAD. This is striking because the conformational change of integrin α_IIb_β_3_ in its active form is a prerequisite for (spontaneous) platelet aggregation^28^. In another study, high glycocalicin levels were detected in patients with AAA, suggesting increased platelet destruction due to elevated platelet activation in these patients. Consequently, the authors detected decreased platelet counts. However, no alterations in platelet counts were observed in the here investigated AAA cohort compared to age-matched controls. Notably, while our study compares platelet counts between patients with AAA and age-matched healthy volunteers, Milne and colleagues used carotid artery stenosis patients as controls^29^. An increase in platelet activation was also observed after endovascular repair of AAA^30^, while our analysis of platelet activation in AAA patients occurred before any treatment.

In addition to enhanced platelet activation in mice and patients, we detected elevated PS exposure at the surface of platelets as a marker for pro-coagulant activity. Enhanced pro-coagulant activity might result from elevated platelet activation. However, it has been shown that inflammatory cytokines may enhance pro-coagulant activity of platelets^31^. Pro-coagulant activity of resting and activated platelets might be responsible for elevated thrombin plasma levels in AAA patients^32,33^ which is a common feature in this disease. Thus, it is not surprising that there is a correlation between elevated pro-coagulant activity of platelets (PS exposure) and aortic diameter expansion in PPE-operated mice (**Figure 5F**). However, the correlation is relatively weak, both in regard to the R and the p value that might be due –at least in part- by the variable results from flow cytometric analysis.

So far, a small number of experimental studies have analyzed the role of platelets in AAA. Touat and colleagues have used a xenograft model of rats to induce aneurysms ^17^. They showed that platelet inhibition by abciximab (which blocks integrin α_IIb_β_3_) limited aortic aneurysm expansion, suggesting that platelet stimulation, and in particular integrin α_IIb_β_3_ activation, may be a promising target in the prevention of AAA progression. To date, platelet responses have been analyzed mainly in the Ang-II infusion model. Liu and colleagues showed that treatment of mice with clopidogrel, an antagonist of the ADP receptor P2Y_12_, prevents Ang-II induced AAA formation by protecting the elastic cell lamina in the mouse aorta. Further, the authors found reduced macrophage infiltration and MMP2 expression after clopidogrel treatment^34^. In a more recent study, Owens and colleagues found reduced rupture of established AAAs and improved survival of mice after clopidogrel treatment^15^. However, all these studies have been done in Ang-II infused mice, an experimental model that has been shown to primarily feature aortic dissection as a precursor to aortic aneurysm formation. In agreement with these results obtained from Ang-II-infused mice, we detected reduced inflammation and reduced elastin fragmentation in platelet-depleted PPE-operated mice, suggesting that platelet-mediated processes play a role in both, aortic dissection and aortic aneurysm formation. However, since platelet activation in experimental PPE mice is comparable to platelet activation in AAA patients including elevated platelet pro-coagulant activity, we believe that the PPE mouse model is a more convenient model to analyze platelet induced effects on AAA formation and progression.

In contrast to the studies mentioned above, a recent study of Liu and colleagues provided evidence for a protective effect of platelets in the Ang-II induced mouse model, in which platelet infusion significantly reduced aortic diameter elevation and inflammatory responses^35^. However, differences in the experimental design and the fact that the Ang-II infusion model is a dissection model and does not reflect alterations in platelet activation as observed in PPE mice and in AAA patients might be responsible for differences in results.

The translation of experimental findings into clinical routine has been an issue for AAA in the past. Several trials for different drug classes have failed to confirm a clinical benefit^36–38^. These disease-specific translational difficulties could have several causal reasons, including limitations in experimental mouse models and the multifactorial character of AAA disease. Researchers have not yet succeeded in identifying a decisive and causal factor for AAA development. In the eyes of the authors, there are multiple reasons why platelets could be a major contributor to AAA development. Platelets represent by far the largest cellular compartment of the blood in terms of quantity, and the presence of an ILT suggests continuous activation in the aneurysmatic abdominal segment. This together with the comparable results in platelet activation, pro-coagulant activity and OPN expression and localization in PPE-operated mice and in patients with AAA presented herein, suggest that the PPE model represents an appropriate model to mimic human AAA pathology.

Interestingly, one study of P-selectin knock-out mice has utilized the PPE mouse model^39^. Genetic deficiency of P-selectin was shown to be protective, as these mice showed attenuated aneurysm formation after infusion with porcine pancreatic elastase. However, the authors claim that P-selectin deficiency resulted in reduced inflammation and aortic wall remodeling according to abdominal aortic tissue samples from day 14 which were stained with either hematoxylin/eosin, Masson’s trichrome or Verhoff’s van Gieson. Moreover, P-selectin is exposed not only at the platelet but also at the endothelial membrane. Thus, the presented effects on inflammation and remodeling cannot be attributed solely to platelets and would need to be verified using techniques other than histology.

OPN is a matricellular cytokine that binds to integrins and CD44 and is elevated in many pathological conditions including AAA^40^. It is synthesized by macrophages, vascular smooth muscles cells (VSMCs) and endothelial cells, and its gene expression is highly upregulated in the aortic wall of AAA patients. Here, we show that platelets enhanced OPN gene expression in macrophages but not in VSMCs or aortic fibroblasts. Platelets contribute to elevated OPN plasma levels through the release of OPN from intracellular stores. OPN plays an important role in inflammatory processes in atherosclerosis, cancer and several chronic inflammatory diseases^41^. OPN is also known for its role in repair and remodeling processes of the vasculature. The proliferation, migration and accumulation of VSMCs and the expression of MMPs via the NF-κB signaling pathway are upregulated by OPN, thereby accelerating the degradation of extracellular matrix in AAA^42,43^. We propose that platelets in-part exert their function in AAA through mediation by OPN, and provide strong evidence for OPN to play a role in the recruitment of platelets to the aortic wall and the ILT, suggesting a vicious cycle of platelet-mediated up-regulation of OPN in the plasma and the aortic wall that allows migration of platelets into aortic tissue, amplifying their detrimental effects on the pathology of AAA. Thus, it is not surprising that OPN knock-out mice show attenuated aneurysm formation after Ang-II infusion^44^. In contrast, the analysis of OPN deficient mice in the PPE model showed no protection. However, the study was performed with a limited number of mice, suggesting that the results have to be confirmed by additional experiments^45^. However, OPN is not the only factor through which platelets can affect AAA, because they also amplify inflammation by up-regulating various cytokines in macrophages and fibroblasts (IL-6), and by inducing the up-regulation of proteins involved in ECM remodeling (MMP9, COL1A1), and can contribute to AAA pathology by elevated platelet activation and pro-coagulant activity. Moreover, platelets contain MMP9 and MMP2 that might be released upon platelet activation in AAA, directly contributing to ECM remodeling^46^.

From a clinical point of view, the question arises as to which AAA patient sub-cohort could benefit most from anti-platelet therapy. Major differences in the efficacy of antiplatelet therapy have been linked to the size of the aneurysm. While patients with small-size aneurysms often did not benefit from anti-platelet therapy or even show elevated risk of bleeding, beneficial effects have been provided for patients with large aneurysms^18,38,47,48^. This might be due to the fact that in patient cohorts including very small AAA, an ILT is seldom present^18^. Enlargement of an AAA is usually associated with the development of an ILT^18^. Based on our results, the ILT or its surface could play a prominent role in the recruitment of platelets. From this, ILT-based endpoints could also become more important for considerations of patient management and anti-platelet therapy in the future. Of note, the clinical implementation of ILT surface measurement to conclude could be comparatively simple to implement but may require further prospective patient studies to evaluate critical endpoints. Considering our findings in summary, the question arises whether an alternative platelet aggregation inhibition, possibly based on GPVI receptor inhibition, might be beneficial. Although our results may advocate for such, further mechanistic insights and data from clinical trials are warranted to substantiate this thesis further.

In summary, own data as well as results from different groups highly support the potential for antithrombotic/antiplatelet strategies in patients to modulate the progression and rupture of AAA. It will be of great importance to identify druggable targets to limit platelet activation and pro-coagulant activity for this purpose.

## Acknowledgments

We thank Martina Spelleken for excellent technical assistance, Sandra Szczutkowski and Christos Dimopoulos for histology and platelet adhesion experiments, and the center for Advanced Imaging (CAi) for technical support.

## Author contributions

MUW, HS and ME designed the study. MUW, JM, KJK, TF, AE, YHR, MC, NP, CB, WI, IK, AMP, and JMS performed experiments. MUW, KJK, MC, and ME analyzed and interpreted data. MUW, KJK and ME wrote the manuscript with all authors providing feedback. MUW, JM, KJK and AE contributed equally to this study.

## Data Availability

The original contributions presented in the study are included in the article, further inquiries can be directed to the corresponding author.

