## Supplemental Data 1 for "Crosstalk of platelets with macrophages and fibroblasts aggravates inflammation, aortic wall stiffening and osteopontin release in abdominal aortic aneurysm"

### 1. Supplementary Methods

**1.1 Cell culture of human cell lines.** Human aortic fibroblasts (hAoF, PromoCell®, #C-12380), human aortic smooth muscle cells (hASMC, PromoCell®, #C-12533) and THP-1 (ATCC, #TIB-202) cell lines were cultured in T-75 flasks in 15 mL of their respective media (hAoF, fibroblast growth medium, Sigma-Aldrich, #C23010; hASMC, DMEM medium Gibco™, #11500416; THP-1, RPMI 1640 medium Gibco™, #21875034). THP-1, hASMC and hAoFs medium including 10% fetal bovine serum (FBS) superior (Sigma-Aldrich, #S0615) and 1% penicillin-streptomycin (Gibco™, #15140122). At 70-80% confluence, cells were split 1:5. For this purpose, adherent hAoF were washed twice with DPBS and dissolved with Trypsin-EDTA (Sigma-Aldrich, #T3924). Trypsinization was stopped by addition of medium, and the cell suspension was transferred to a falcon tube and centrifuged at 200 *g* for 5 min. The supernatant was discarded and the cells were resuspended in fresh medium. Non-adherent THP-1 cells do not require trypsinization and were immediately transferred into a falcon tube. Before resuspension in fresh medium the cells were washed once with DPBS by an additional centrifugation step at 200 *g* for 5 minutes. Once resuspended in fresh medium, the cells were ready for cell culture experiments.

For cell culture experiments, cells were seeded into a 24-well plate at 1.5 x $10^5$ cells/well. HAoF were incubated for 24 h at 37 °C and 5% $CO_2$ to allow the cells to become adherent. THP-1 cells were differentiated into macrophages as described elsewhere (1). Therefore, THP-1 cells were differentiated into macrophages by incubation with 150 nM phorbol 12-myristate 13-acetate (PMA, Sigma, #P8139) for 24 hours. After another 24-hour incubation with RPMI medium, the macrophages were polarized into M1 macrophages with 20 ng/ml IFN-γ (R&D System, #285-IF) and 100 ng/ml LPS (Sigma, #8630). These cells were called “differentiated macrophages”. Once hAoF and macrophages became adherent, the platelet supernatant was added and the cells were incubated at 37 °C and 5% $CO_2$ for 6 h. Afterwards, the supernatant was discarded and the cells were washed twice with ice-cold DPBS. Finally, the cells were detached from wells using TRI Reagent® (Sigma-Aldrich, #T9424).

**1.2 Gene expression from human platelets and cell lysates of macrophages and hAoFs.** Human platelets were isolated from platelet concentrates using the following protocol. 1-5 x $10^9$ of either resting or activated platelets were centrifuged at 2500 *g* for 5 min at 4 °C. The supernatant was discarded and the pellet was resuspended in TRI Reagent®. After transferring the sample into a DNA LoBind Tube (Eppendorf, #30108051), Chloroform (Sigma-Aldrich, #372978) was added at a ratio of 1:5. The samples were shaken thoroughly, incubated for 3 min at RT and centrifuged at 12,000 *g* for 15 min at 4 °C. The aqueous phase was diluted in pure ethanol (3-fold) and mixed with linear acrylamide (1:500) (Invitrogen, #AM9520). After an incubation at 4 °C overnight, the samples were centrifuged at 21,130 *g* at 4 °C. The supernatant was discarded, and the pellet was washed in ice-cold 80% ethanol and resuspended in 25 µL nuclease free water. Total RNA from cell culture lysates of differentiated macrophages and hAoFs was isolated using the RNeasy Mini Kit (Qiagen, #74106) according to the manufacturer's protocol. RNA content and quality were measured using the BioPhotometer D30 (Eppendorf). All samples were adapted to the same amount of RNA. DNA digestion was performed with DNAse I (Roche, #04716728001) according to the following protocol: 30min/37 °C; 10min/75 °C using a PCR thermocycler (Eppendorf). Subsequent cDNA synthesis was performed using the ImProm- II™ Reverse Transcription System (Promega, #A3800) according to their protocol. For quantification of mRNA levels, fast SYBR™ Green Master Mix (Applied Biosystems, #4385612) was used with following specific oligonucleotide primers: MMP2 (PPH00151B-200), MMP9 (PPH00152E-200), COL1A1 (PPH01299F-200), COL3A1 (PPH00439F-200), SPP1 (PPH00582E-200), and GAPDH (PPH00150F-200) acquired from Qiagen. Following oligonucleotide primers were acquired from Origene: MCP1 (HP 206587), IL-6 (HP 200567), IL-12β (HP 205923), TNF-α (HP 200561), IL-10 (HP 200540) and IL-8 (HP 200551). The IL-1β primer pair (frwd: AGCTTCCTTGTGCAAGTGTCTGAG, rev: TGTTGATGTGCTGCTGCGAGAT) and IL-6 (hAoFs) primer pair (frwd: GGCACTGGCAGAAAACAACC; rev. GCAAGTCTCCTCATTGAATCC) were purchased from Eurofins. Data are normalized to GAPDH and fold changes were calculated using the $^{\Delta\Delta}$Ct method.

**1.3 Gene expression of murine aortic tissue.** Total RNA was isolated from murine aortae with a TRIzol-based (Invitrogen) RNA isolation protocol. RNA was quantified by NanoDrop (Agilent Technologies), and RNA and miRNA quality were verified with the Agilent 2100 Bioanalyzer (Agilent Technologies). Samples required 260/280 ratios of >1.8 and sample RNA integrity numbers of ≥9 for inclusion. RNA was reverse-transcribed with the TaqMan MicroRNA Reverse Transcription kit (Applied Biosystems) according to the manufacturer's instructions. To quantitate the mRNA levels the TaqMan qRT-PCR assay was used. The following specific oligonucleotide primers were purchased from Thermo Fisher Scientific:

*Il-1β* (Mm00434228_m1), *Il-6* (Mm00446190_m1), *Il-8* (Mm00441263_m1), *Il-10* (Mm01288386_m1), *Il-12β* (Mm01288989_m1), *Ccl2* (Mm00441242_m1), *Tnf-α* (Mm00443258_m1), *Eln* (Hs00355783_m1), *Mmp2* (Mm00439498_m1), *Mmp9* (Mm00442991_m1), *Col1a1* (Mm00801666_g1), *Col3a1* (Mm01254476_m1), *Spp1* (Mm00436767_m1), *18S* (4319413E). Data were normalized to 18S or GAPDH and all fold changes were calculated using the $^{\Delta\Delta}$Ct method.

**1.4 Fragmentation of osteopontin.** Fragmentation of recombinant osteopontin (OPN) (PeproTech, #120-35) was performed with thrombin (30 mU/1 µg OPN) (Roche, #10602400001) by incubation at 37 °C for 3 h. Silver staining was used to verify the concentration of enzymes and the quality of osteopontin fragmentation. Therefore the product was separated via gel electrophoresis and stained using Pierce Silver Stain Kit (Thermo Fisher Scientific, #24612) according to the manufacturer's protocol.

**1.5 Adhesion studies.** Adhesion experiments were performed to investigate adhesion of human platelets on various protein matrices. For the immobilized osteopontin experiments, full-length osteopontin (fl. OPN, 5 µg/mL) (PeproTech, #120-35) and cleaved osteopontin (cle. OPN, 5 µg/mL) were applied to glass coverslips (24 x 60 mm) in a defined area (10 x 10 mm). For the experiments with soluble osteopontin, the glass coverslips were coated with collagen (200 µg/mL) (Horm®, Takeda Pharmaceutical, #3408075). A matrix of bovine serum albumin (BSA, 1%) was used as negative control and fibrinogen (100 µg/mL) (Sigma-Aldrich, #F3879) was used as a positive control. To generated coated surfaces, glass coverslips were incubated in a humidity chamber at 4 °C overnight and blocked with 1% BSA at RT for 1 h. Human platelets were isolated from the blood of healthy volunteers and treated as required in the following experiments. For the experiments with immobilized osteopontin, platelets were pre-treated with or without an inhibiting $\alpha_v\beta_3$-antibody (30 µg/mL) (Sigma-Aldrich, #MAB1976) at 37 °C for 15 min. Resting, ADP (10 µM; Sigma-Aldrich, Germany, #A2754) and CRP (1 µg/mL; collagen-related peptide, CambCol Laboratories, United Kingdom, #28045457) stimulated platelets (40,000/ µL) were added to the matrices and incubated at 37 °C for 30 min. For the soluble osteopontin experiments, platelets were pre-treated with fl. OPN (1 µg/mL) and cle. OPN (1 µg/mL) at 37 °C for 30 min and then added to a collagen matrix for 5 or 20 min. Finally, unbound platelets were washed off with DPBS, and adhesive platelets were fixed with ice-cold 4% paraformaldehyde for 10 min. Permeabilization with Triton-X100 (0.5%) (Sigma-Aldrich, #T8787) in BSA solution (0.5%) for 10 minutes served as preparation for actin filament staining with rhodamine-phalloidin (1:200) (Invitrogen, #R415) for 40 min. Subsequently, the cover slips were washed with DPBS and covered with the mounting medium Fluoromount-G™ (Sigma-Aldrich, #F4680) and monitored by microscopic. Five pictures from different areas of one biological replicate were taken (platelets: 400 x total magnification; Axio Observer.D1, Carl Zeiss). The total number of adherent platelets was counted using ZEN 3.3. (blue edition) software.

Spreading experiments were performed as described previously (2-4). For spreading analysis of isolated murine platelets the adherent platelets were rinsed, fixed and covered as described above. The samples were imaged using Microscope Axio Observer.D1 (Objective Plan Apochromat 100x/1.40 Oil DIC M27) for differential interference contrast (DIC) imaging resulting in at least 5 representative visual field pictures per slide for 1 sample per mouse. The whole visual field was counted by marking the different spreading stages in Carl Zeiss Software ZEN 3.3 (blue edition). Platelets that show only filopodia were counted as filopodia forming cells, platelets that start to form lamellipodia and fully spread platelets were counted as lamellipodia forming platelets and landing and attached platelets were counted as adherent cells according to Aslan and McCarty, 2012 (5). Users were blinded to treatment.

For immunofluorescence staining of OPN on adherent platelets, 4.9 x $10^5$ platelets were placed on matrices coated with collagen and fibrinogen. After fixation with 2% paraformaldehyde and permeabilization of platelets, nonspecific binding was blocked with goat serum (5%) (Thermo Fisher Scientific, #01-6201) in Tris-buffered saline with 0.1% Tween® 20 detergent (TBS-T) buffer for 1 h. The primary antibodies (20 µg/mL) (abcam, OPN #ab181440, rb IgG #ab27478) were incubated at 4 °C overnight. Secondary antibodies (Invitrogen, 20 ng/mL, goat anti-rabbit 488 #A11008; 3U rhodamine-phalloidin) were incubated at RT for 1 hour. After the staining and mounting with Fluoromount-G™, images were captured using a Leica DMi8 confocal microscope.

**1.6 Thrombus formation under flow using the *ex vivo* flow chamber system.** Glass cover slips (24 x 60 mm) were coated with collagen (200 µg/mL) or a combination of collagen (200 µg/mL) and fl. OPN (100 µg/ml) (Peprotech, #120-35) at 4 °C overnight. On the next day, spare collagen was removed and cover slips were blocked with BSA (1%) at RT for at least 1 hour. Human whole blood was perfused over both matrices through the flow chamber (diameter: 50 µm x 5 mm) at a shear rate of 1,700 $s^{-1}$ using a pulse-free electric pump. After 3 min, blood perfusion was stopped and Tyrode's buffer was pumped through the flow chamber using the same shear rate. Five pictures were taken (400 x total magnification; Axio Observer.D1, Carl Zeiss) from different areas. The surface coverage was analyzed using ImageJ-win64 software.

**1.7 Histology.** For the analysis murine or human aortic tissue (aortic wall), 5-7-micron sections of paraffin-embedded tissue were prepared. Additionally, cryopreserved human thrombus samples (intraluminal thrombus, ILT) were prepared by embedding the tissue into Tissue-Tek® O.C.T.™ compound. Sections were cut using the Leica Biosystems (CM1950 Cryostat). Hematoxylin and eosin staining was performed using a common HE-staining protocol (2). For quantification of the intima/media thickness 2 fields per cross section were analysed by measuring media thickness at 10 distinct locations within each cross section using the ZEN 3.3. (blue edition) software. The modified Elastin Verhoeff van Gieson staining (VVG) was conducted according to manufacturer's protocol (abcam, elastic stain kit #ab150667). Elastin fragmentation was determined as elastin fragments / media lamellar units via ZEN 3.3. (blue edition) software. The number of these breaks were counted in both thoracic and abdominal aortic segments of naive mice, platelet-depleted PPE-operated mice, and IgG treated PPE mice (controls). For each tissue sample 2 fields per cross section were analyzed.

**1.8 Immunofluorescence staining.** For immunofluorescence staining of osteopontin (OPN), platelets (CD42b), macrophages (Mac-3) and fibroblasts (Vimentin), sections of human and murine aortic walls as well as intraluminal thrombi (ILT) of AAA patients were stained. For staining of human and murine aortic wall, paraffin-embedded tissue was used, while staining of human ILT was performed in cryopreserved tissue. Prior to the staining, the paraffin-embedded sections were deparaffinized and hydrated. For antigen unmasking the tissues were microwaved at 360 kW for 10 min. ILT sections were fixed with 4% paraformaldehyde for 10 min before staining. For the blocking of nonspecific binding sites, sections of both, cryopreserved and paraffin-embedded were incubated with DPBS + 0.3 % Triton x-100 and goat serum (5%) at RT for 1 h. The primary antibodies ( (anti-osteopontin, 20 µg/mL, abcam, #ab181440), recombinant rabbit IgG (monoclonal [EPR25A] isotype control (20 µg/mL, abcam, #ab172730)), CD42b monoclonal antibody (20 µg/mL, Invitrogen™, #42C01), BD Pharmingen™ purified mouse IgG1 κ isotype control (50 µg/mL, BD Bioscience, #554121), BD Pharmingen™ purified rat anti-mouse Mac-3, unlabeled (3.125 µg/mL, BD Bioscience, #BDB550292), anti-CD68 AA 21-120 (10 µg/mL, antibodies-online GmbH, #ABIN671406), anti-CD68 (10 µg/mL, antibodies-online GmbH, #ABIN6941238), vimentin polyclonal antibody (20 µg/mL, Invitrogen™, #PA5-142829), purified rat anti-mouse GPIbα (CD42b, 10 µg/mL, Emfret Analytics, #M042-0)) were incubated at 4 °C overnight. On the next day, the appropriate secondary antibodies (goat anti-rabbit IgG (H+L) cross-adsorbed secondary antibody, Alexa Fluor™ 555 (20 µg/mL, Invitrogen™, #A-21428), eBioscience™ streptavidin eFluor™ 660 conjugate (10 µg/mL, Invitrogen™, #50-4317-80)) were applied to the tissues at RT for 1 h. Negative controls were generated by omission of the primary antibody. In case of platelet staining with CD42b, an intermediate step of biotinylation was used to enhance the fluorescence signal (goat anti-mouse IgG antibody (H+L), biotinylated, GT (7.5 µg/mL, Vector Laboratories, #VEC- BA-9200), goat anti-rat IgG antibody (H+L), biotinylated (7.5 µg/mL, Vector Laboratories, #BA-9400)). Nuclei were identified using DNA staining with 4, 6 diamidino-2-phenylindole dihydrochloride (DAPI, Roche, 1:3000). Samples were analysed using confocal microscopy (LSM 710, Carl Zeiss, Germany). For quantification of the platelet, macrophage and OPN content within the aortic tissue of PPE-operated mice the mean fluorescence intensity (MFI) of 2 subsequent sections was used. For every cross-section 2 representative fields were analysed. The MFI of specifically stained sections was automatically normalized to the area of total aortic tissue. Determination of the normalized MFI was performed using the Carl Zeiss Software ZEN 3.3 (blue edition). For image generation and quantification users were blinded to the treatment.

**1.9 Quantification of plasma levels of OPN, MMP2, MMP9 and IL-6.** Circulating OPN in plasma of mice was determined according the manufacturer's protocol using mouse/rat osteopontin (OPN) Quantikine ELISA Kit (R&D Systems, #MOST00). In addition, OPN content was analyzed in human plasma, isolated human platelets (stimulated with 10 µM ADP), and from differentiated macrophage cultures using the osteopontin/SPP1 ELISA kit (Thermo Fisher

Scientific, #EHSPP1) according to manufacturer specified protocol. IL6 concentration of hAoF cultures were determined using the Human IL-6 DuoSet ELISA kit (R&D Systems, #DY206-05) according to the manufacturer protocol. MMP2 (R&D Systems, #DY902) and MMP9 (R&D Systems, #DY911) levels in human plasma were also determined according to the manufacturer's protocol.

**1.10 Experimental animals: The Ang-II infused ApoE knockout mouse model.**

For angiotensin-II (Ang-II) induced AAA formation, 10-week-old male *ApoE*$^{-/-}$ mice purchased form the Jackson Laboratory were used. Prior implantation the alzet osmotic mini-pumps (model 1004, Durect Corporation) were filed with Ang-II (Sigma-Aldrich, #A92525) or sterile isotonic saline (NaCl, 0.9%) and were primed at 37°C in sterile 0.9% $NaCl_2$ overnight. For the implantation mice were anesthetized with 2-3% isoflurane and additionally received a locally subcutaneous (s.c.) injection of metamizol (200 mg/kg) 30 min before surgery. The osmotic mini-pumps were implanted into a subcutaneous pocket within the dorsal region caudal to the scapula. After implantation the osmotic mini-pumps reach a delivery rate of 1 µg/kg/min over a time period of 28 days. For pain relief, all animals received metamizol via drinking water (1.33 mg/mL) for two days. For platelet depletion Ang-II infused *ApoE*$^{-/-}$ mice received a platelet depletion antibody (Emfret Analytics, polyclonal anti-GPIb alpha #R300) or a corresponding IgG control antibody (Emfret Analytics, #C301) after surgery (days 0 and 5). The mice were sacrificed at day 7 post osmotic mini-pump implantation.

**1.11 Ultrasound imaging.** Prior either PPE-surgery or osmotic mini-pump implantation (baseline) and at days 3, 7, 10, 14, 21 and 28 following surgery, maximal aortic diameters at the aneurysm site were measured using ultrasound. Mice were anaesthetized with 2-3% isoflurane, placed on a 37 °C heated plate and ultrasound imaging was performed using a Vevo 2100® High-Resolution In Vivo Micro-Imaging System (VisualSonics). Inner diameter-measurements were obtained following a standardized imaging algorithm with longitudinal B-Mode images during the systolic phase. PWV was examined to assess aortic stiffness by simultaneous tracking of the ECG and the pulse wave at the aortic bifurcation (bif). The distance from the aortic valve to the bif was measured during sacrifice and was set to 3.98 cm for further calculations. For each mouse, 5 PW Doppler sets were recorded. In each set, 3 measurements from the foot of the R-wave of the ECG to the offspring of the pulse wave at the bif were measured to determine the transit time between these two locations. Global aortic PWV was estimated as a ratio of the distance (d) and time (t) delay of the pulse wave between both locations.

### 2. Supplemental Figures

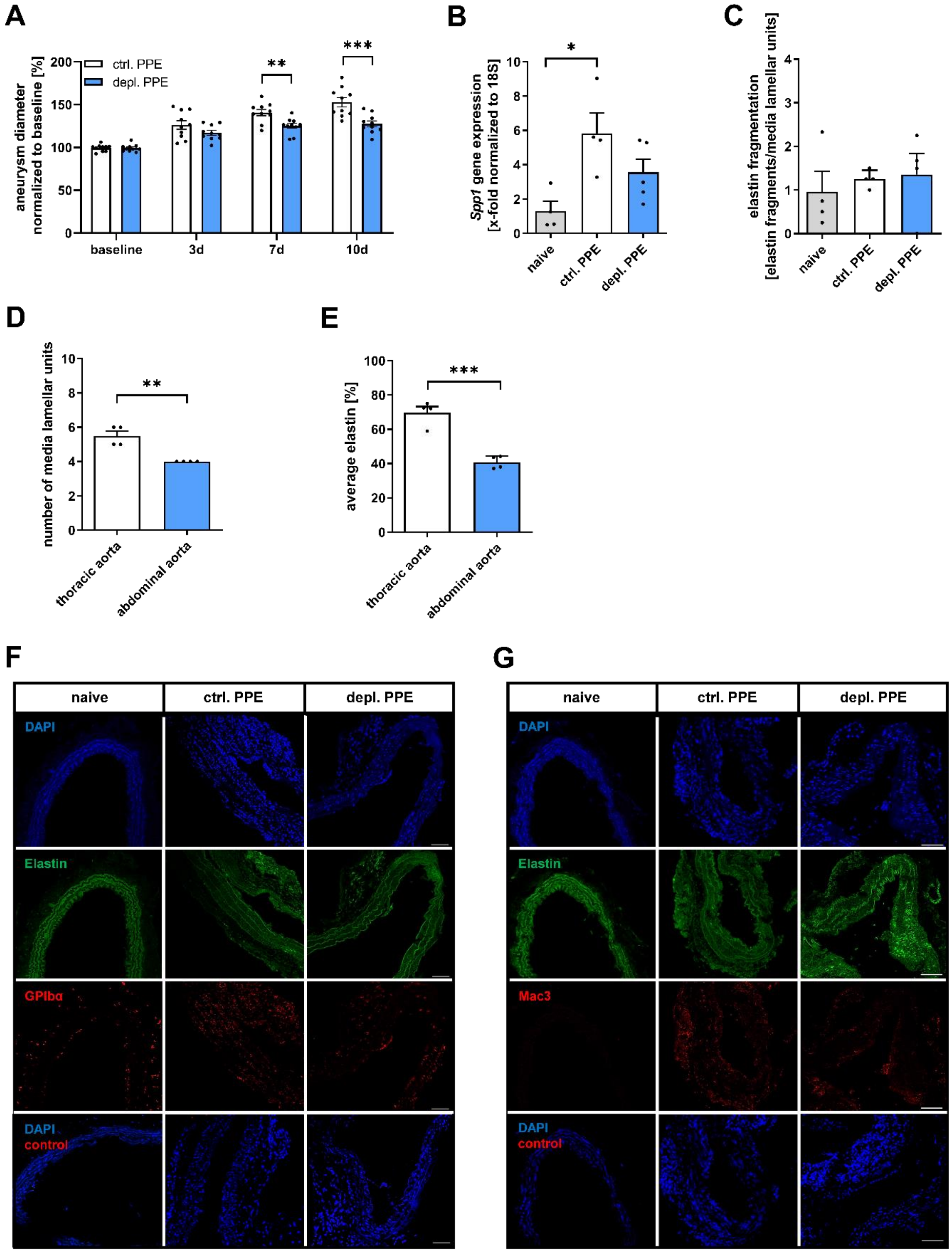


**Figure S1. Platelet depletion reveals unaltered elastin degradation within the thoracic aorta in the PPE mouse model.** (**A**) Aneurysm diameter of depl. PPE and ctrl. PPE mice was determined by ultrasound measurements at the indicated time points (n = 10). Data were normalized to baseline. (**B**–**E**) Different tissue analysis of thoracic aortae depl. PPE and ctrl. PPE mice. (**B**) *Spp1* gene expression in thoracic aortic tissue (n = 4–5). (**C**) Elastin fragmentation counted in Verhoeff van Gieson-stained samples from naive and PPE-operated mice (n = 4). (**D**) Number of lamellar units within the thoracic and abdominal aorta segment of naive mice (n = 4). (**E**) Average elastin content of thoracic and abdominal aortic tissue of naive mice as a percentage of whole depicted tissue amount in the sample (n = 4). (**F** and **G**) Representative immunofluorescence (IF) images of (**F**) platelets (CD42b/anti-GPIbα, red) or (**G**) macrophages (anti-Mac3, red), nuclei (DAPI, blue) and elastin (autofluorescence, green) in aortic tissue sections of depl. PPE and ctrl. PPE at day 3 post-surgery (n = 3–5). Scale bar: 50 µm. Aortic tissue of naive mice served as control (n = 4). Negative controls were achieved by using the respective IgG antibody (control). Data are represented as mean ± SEM. Statistical significances were determined by two-way ANOVA with Sidak's multiple comparison test (**A**), one-way ANOVA with Holm Sidak's multiple comparisons test (**B** and **C**) or unpaired student's t-test (**D** and **E**). *$P < 0.05$, **$P < 0.01$, ***$P < 0.001$. Ctrl., control; depl., depletion; PPE, porcine pancreatic elastase (infusion); PWV, pulse wave velocity; *Spp1*, secreted phosphoprotein 1.

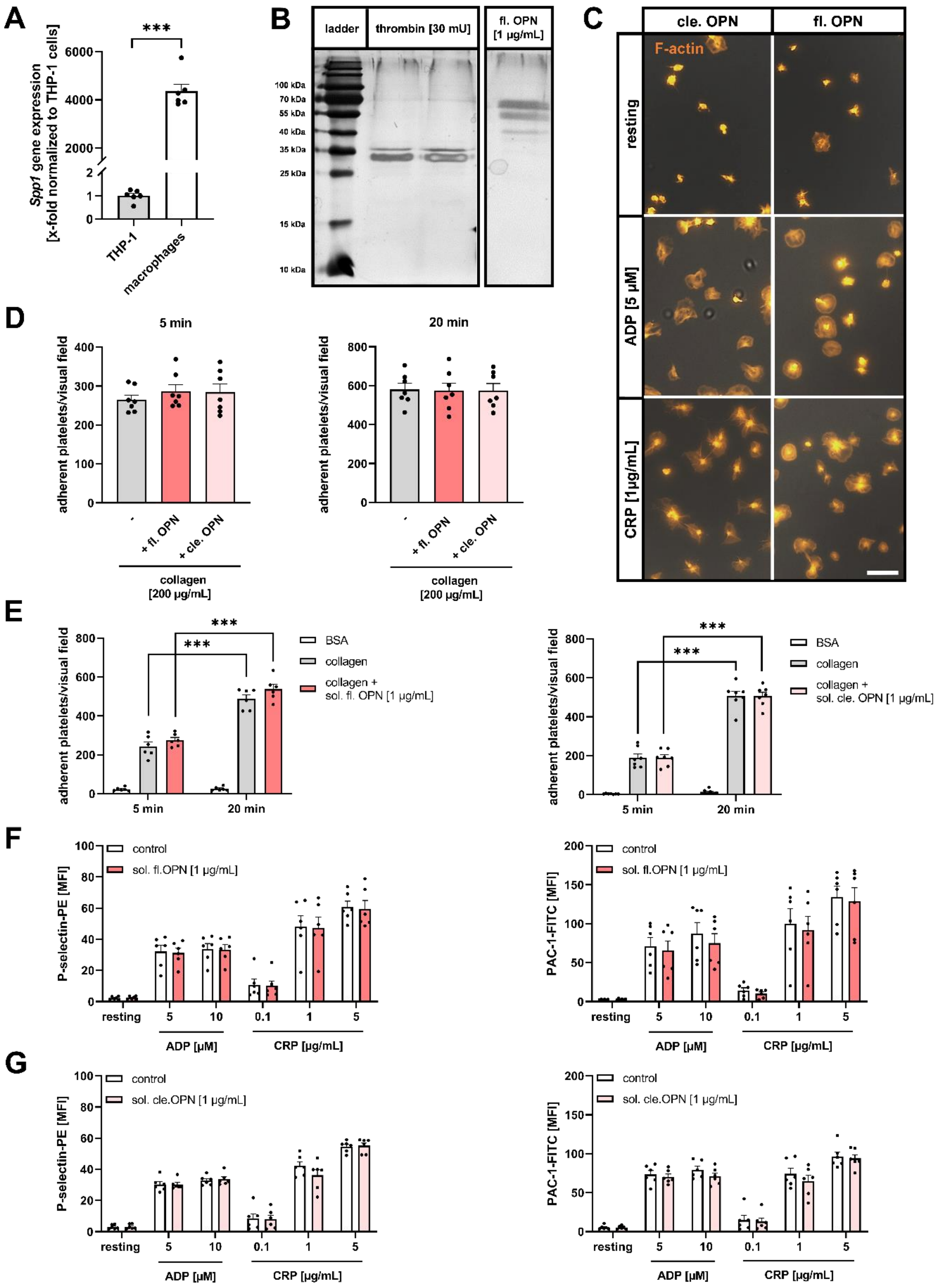

**Figure S2. OPN does not modulate adhesion to collagen and platelet activation under static conditions.** (**A**) *SPP1* gene expression in THP-1 cells and differentiated macrophages (n = 6). Fold changes were normalized to THP-1 *SPP1* gene expression. (**B**) Full-length OPN was cleaved by thrombin (30 mU/ 1 µg OPN). Silver staining was used to verify the quality of OPN fragmentation. The products were separated via gel electrophoresis and stained using a Pierce Silver Stain Kit. (**C**)

Representative images of filopodia- and lamellipodia-forming platelets spread on cleaved or full-length osteopontin (5 µg/mL) for 30 minutes. Platelets were stimulated with indicated agonists. Adherent platelets were visualized by rhodamine-phalloidin staining (F-actin, orange) (n = 5). Scale bar: 10 µm. (**D**) Isolated platelets were allowed to adhere on a collagen (200 µg/mL) or collagen/OPN (full-length or cleaved, 5 µg/mL) matrix for 5 and 20 min (n = 7). (**E**) Quantification of adherent human platelets, pre-treated with soluble full-length (fl. OPN, 1 µg/mL) or cleaved (cle. OPN, 1 µg/mL) OPN on a collagen matrix after 5 and 20 min (n = 6–7). BSA served as control. (**F** and **G**) Platelet degranulation (P-selectin-PE) and active integrin $\alpha_{IIb}\beta_3$ externalisation (PAC-1-FITC) of human platelets after incubation with or without (control) full-length or cleaved OPN (1 µg/mL) were determined by flow cytometry (n = 6). Platelets were stimulated with indicated agonists. Data are represented as mean ± SEM. Statistical significances were determined by unpaired student‘s t-test (**A**), one-way ANOVA with Holm-Sidak’s multiple comparisons test (**D**), two-way ANOVA with Tukey’s multiple comparisons test (**E**) or multiple t-test (**F** and **G**). ***$P < 0.001$. ADP, adenosine diphosphate; BSA, bovine serum albumin; cle., cleaved; CRP, collagen-related peptide; fl., full-length; OPN, osteopontin; sol., soluble.

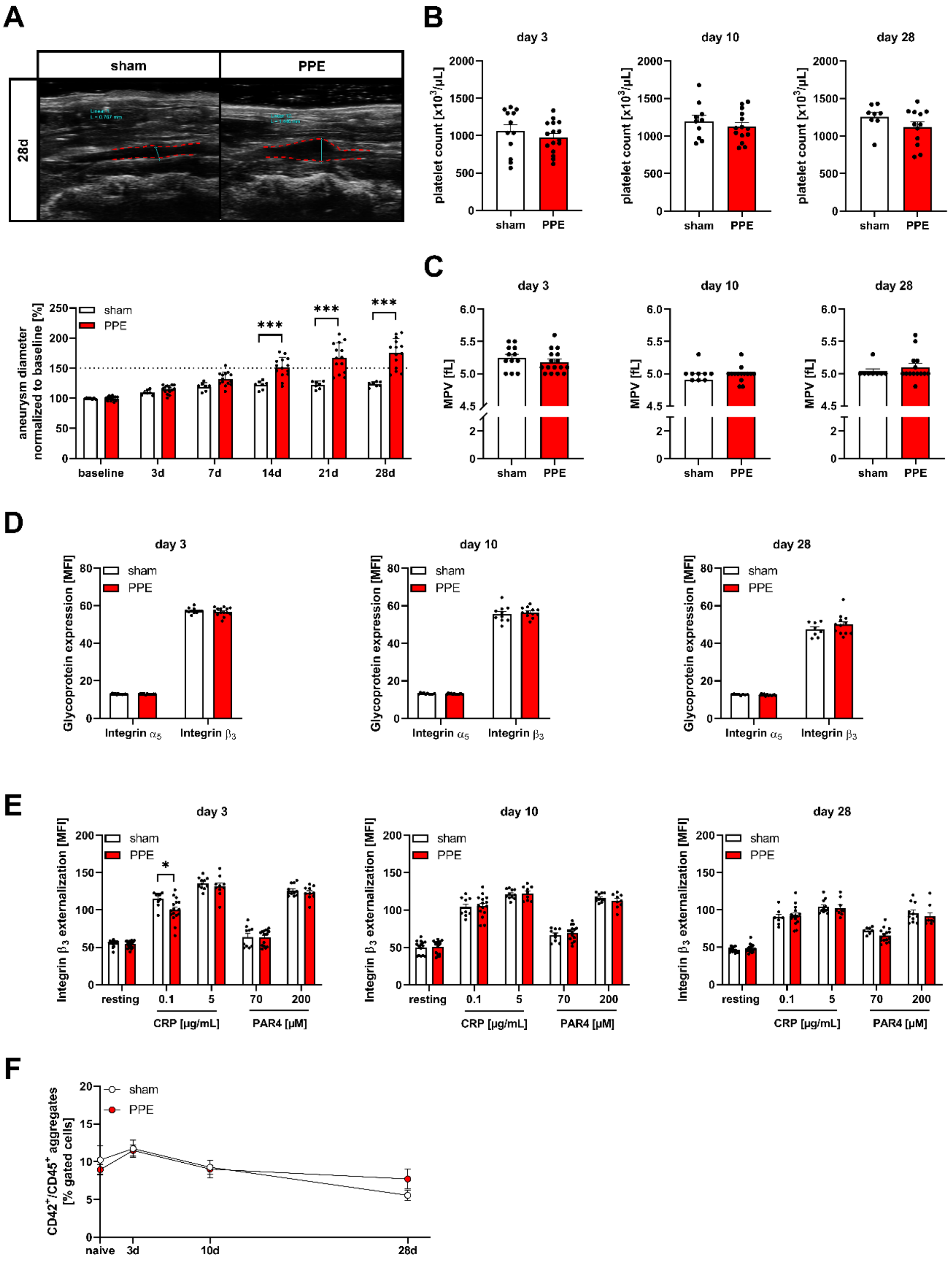


**Figure S3. Platelet count, MPV and glycoprotein externalization are not influenced in PPE-operated mice.** (**A**) Representative ultrasound images (at day 28) and aortic diameter progression of sham respectively PPE-operated mice over a time-period of 28 days after surgery (n = 8–13). Data were normalized to baseline. (**B**) Platelet count of sham and PPE-operated mice at days 3, 10 and 28 post-surgery (n = 8–16). (**C**) MPV of sham and PPE-operated mice 3, 10 and 28 days post-surgery (n = 8–16). (**D**) Externalization of integrins $\alpha_5$ and $\beta_3$ on the platelet surface in whole blood from mice at days 3, 10 and 28 after sham or PPE surgery (n = 10–12). (**E**) Externalization of integrin $\beta_3$ on the platelet surface in whole blood from mice at days 3, 10 and 28 after sham or PPE surgery after stimulation with indicated agonists (n = 10–12). (**F**) Aggregate formation of platelets ($CD42^+$) and leukocytes ($CD45^+$) in whole blood of naive, sham and PPE-operated mice at days 3, 10 or 28 post-surgery determined by flow cytometry (n = 5–6). Data are represented as mean or percent-gated cells ± SEM. Statistical significances were determined by two-way ANOVA with Sidak's multiple comparison test (**A**), unpaired student's t-test (**B** and **C**), multiple t-test (**D** and **E**) or two-way ANOVA with Sidak's multiple comparison test (**F**). $^*P < 0.05$. CRP, collagen-related peptide; MFI, mean fluorescence intensity; MPV, mean platelet volume; PAR4, protease-activated receptor 4 activating peptide; PPE, porcine pancreatic elastase (infusion).

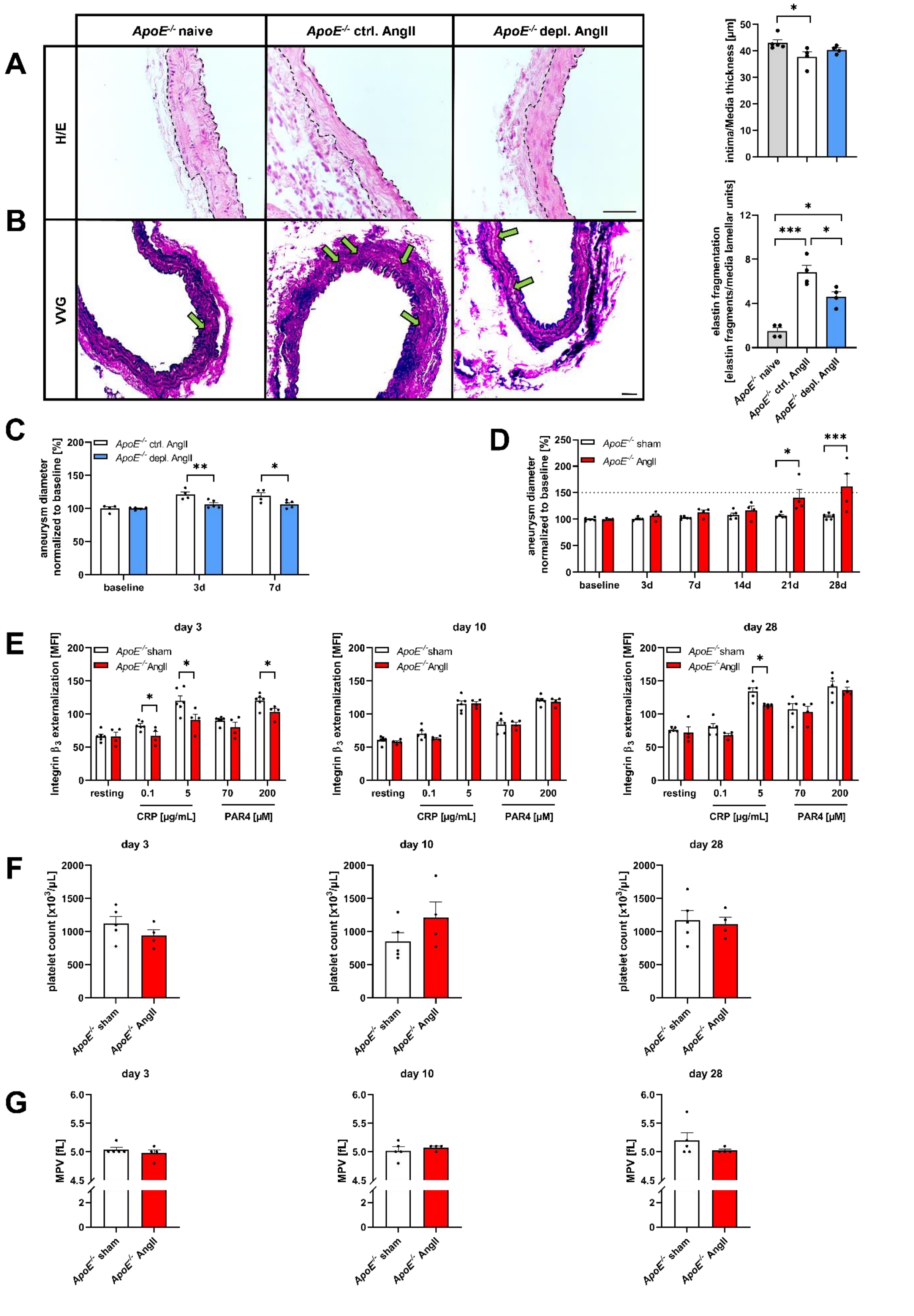


**Figure S4. Aortic remodeling, platelet integrin β$_3$ externalization, platelet counts and MPV in ApoE knockout mice with experimental angiotensin-II induced AAA formation.** (**A** and **B**)

Representative images of aortic tissue from depl. *ApoE*$^{-/-}$ and ctrl. *ApoE*$^{-/-}$ mice treated with Ang-II analysed via (**A**) hematoxylin-eosin (H/E) staining and (**B**) Verhoeff van Gieson (VVG) staining. Intima/media thickness was determined after H/E staining (n = 4–5) and elastin fragmentation (breakdowns) was counted after VVG staining (n = 4). Aortic tissue of naïve mice served as control. Scale bars: (**C** and **D**) 50 µm. (**C**) Aortic diameter progression of depl. *ApoE*$^{-/-}$ and ctrl. *ApoE*$^{-/-}$ mice infused with Ang-II for 7 days measured via ultrasound. Data were normalized to baseline (n = 4–5). (**D**) Aortic diameter progression of sham or Ang-II infused *ApoE*$^{-/-}$ mice over a time-period of 28 days determined by ultrasound measurements (n = 4–5). Data were normalized to baseline. (**E**) Externalization of integrin $\beta_3$ on the platelet surface in whole blood from mice at days 3, 10 and 28 of sham or Ang-II infused *ApoE*$^{-/-}$ mice after stimulation with indicated agonists (n = 4–5) (**F**) Platelet count of sham or Ang-II infused *ApoE*$^{-/-}$ mice at days 3, 10 and 28 (n = 4–5). (**G**) MPV of sham or Ang-II infused *ApoE*$^{-/-}$ mice 3, 10 and 28 days after osmotic mini-pump implantation (n = 4–5). Data are represented as mean ± SEM. Statistical significances were determined by one-way ANOVA with Holm-Sidak's multiple comparison test (**A** and **B),** two-way ANOVA with Sidak's multiple comparison test (**C** and **D**), multiple t-test (**E**) or unpaired student's t-test (**F** and **G**) or $^*P < 0.05$, $^{**}P < 0.01$, $^{***}P < 0.001$. Ang-II, angiotensin-II; CRP, collagen-related peptide; ctrl, control; depl., depletion; H/E, hematoxylin-eosin; MFI, mean fluorescence intensity; MPV, mean platelet volume; PAR4, protease-activated receptor 4 activating peptide; VVG, Verhoeff van Gieson.

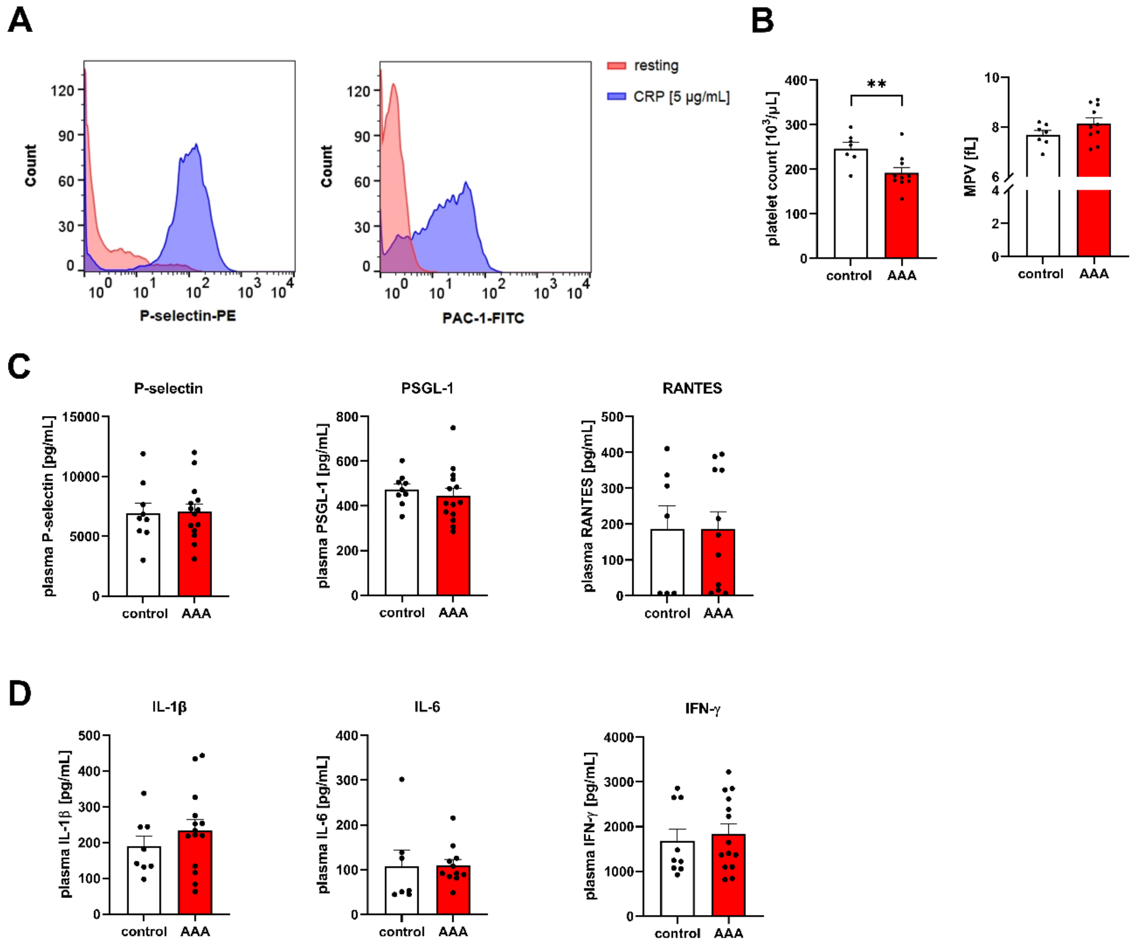


**Figure S5. Platelet count, MPV and plasma parameters of AAA patients.**

(**A**) Representative histograms for the flow cytometric analysis of P-selectin (left panel) and PAC-1 (right panel) of platelets from AAA patients under resting conditions and after stimulation with 5 µg/mL CRP. (**B**) Platelet count and MPV in AAA patients and age-matched controls (n = 6–10). (**C**) Plasma concentration of P-selectin, PSGL-1 and RANTES in AAA patients and age-matched controls determined by flow cytometric multiplex analysis. (**D**) Plasma concentration of IL-1β, IL6 and IFN-γ in samples of AAA patients and age-matched controls determined by flow cytometric multiplex analysis (n = 7–14). Data are represented as mean ± SEM. Statistical significances were determined by unpaired t-test with Welch's correction (**B**–**D**). ***P* < 0.01. AAA, abdominal aortic aneurysm; CRP, collagen-related peptide; IFN, interferon; IL, interleukin; MPV, mean platelet volume; PSGL, P-selectin glycoprotein ligand; RANTES, regulated on activation, normal T cell expressed and secreted.

**A**

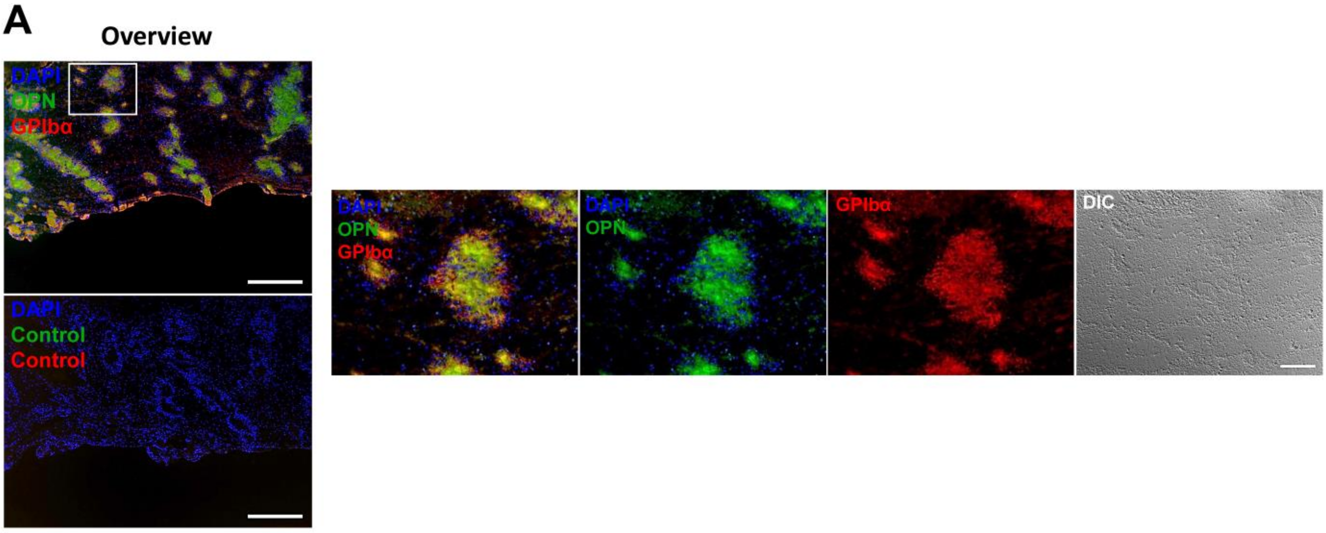
Overview
DAPI
OPN
GPIbα
DAPI
Control
Control
DAPI
OPN
GPIbα
DAPI
OPN
GPIbα
DIC

**B**

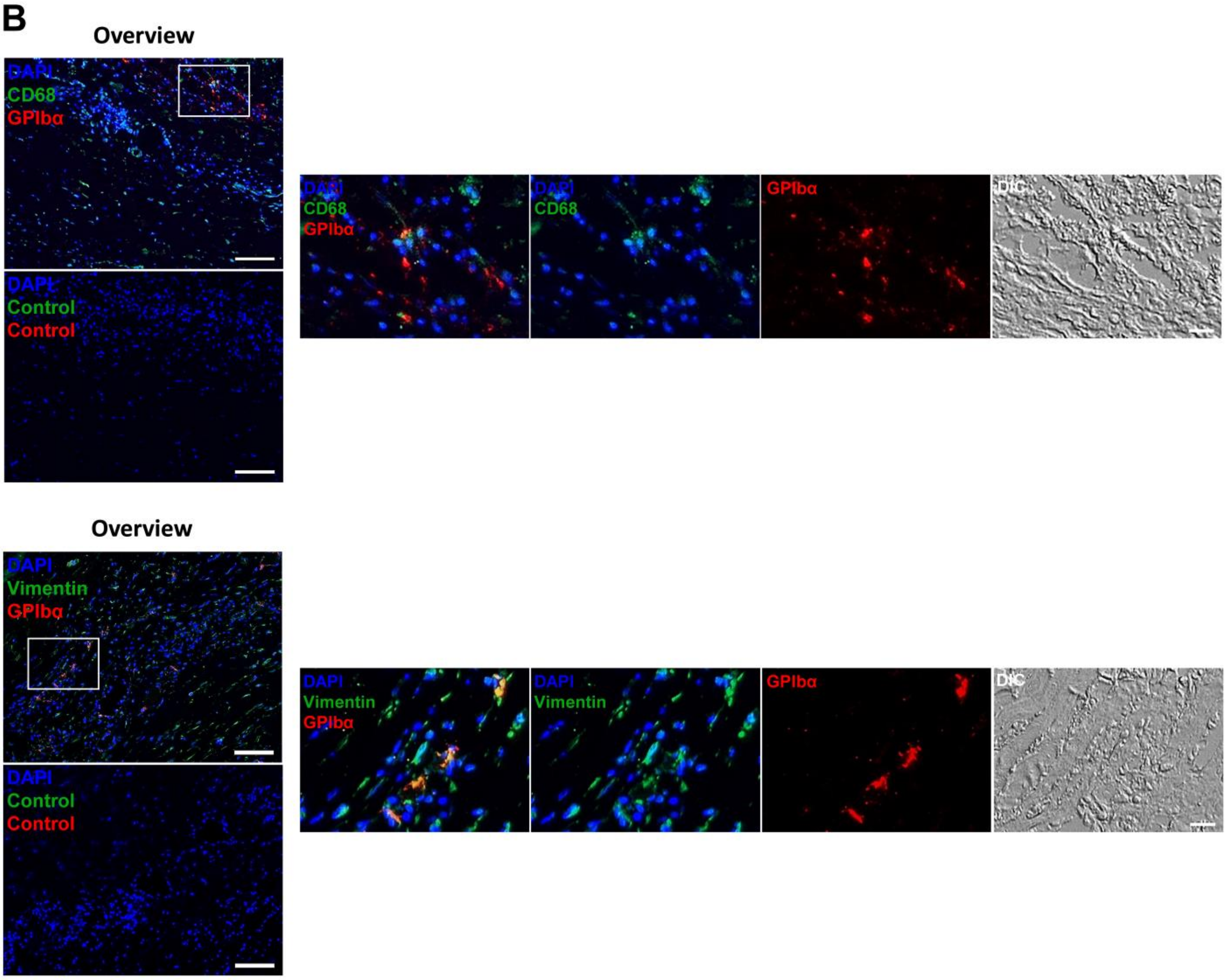
Overview
DAPI
CD68
GPIbα
DAPI
Control
Control
DAPI
CD68
GPIbα
DAPI
CD68
GPIbα
DIC
Overview
DAPI
Vimentin
GPIbα
DAPI
Control
Control
DAPI
Vimentin
GPIbα
DAPI
Vimentin
GPIbα
DIC

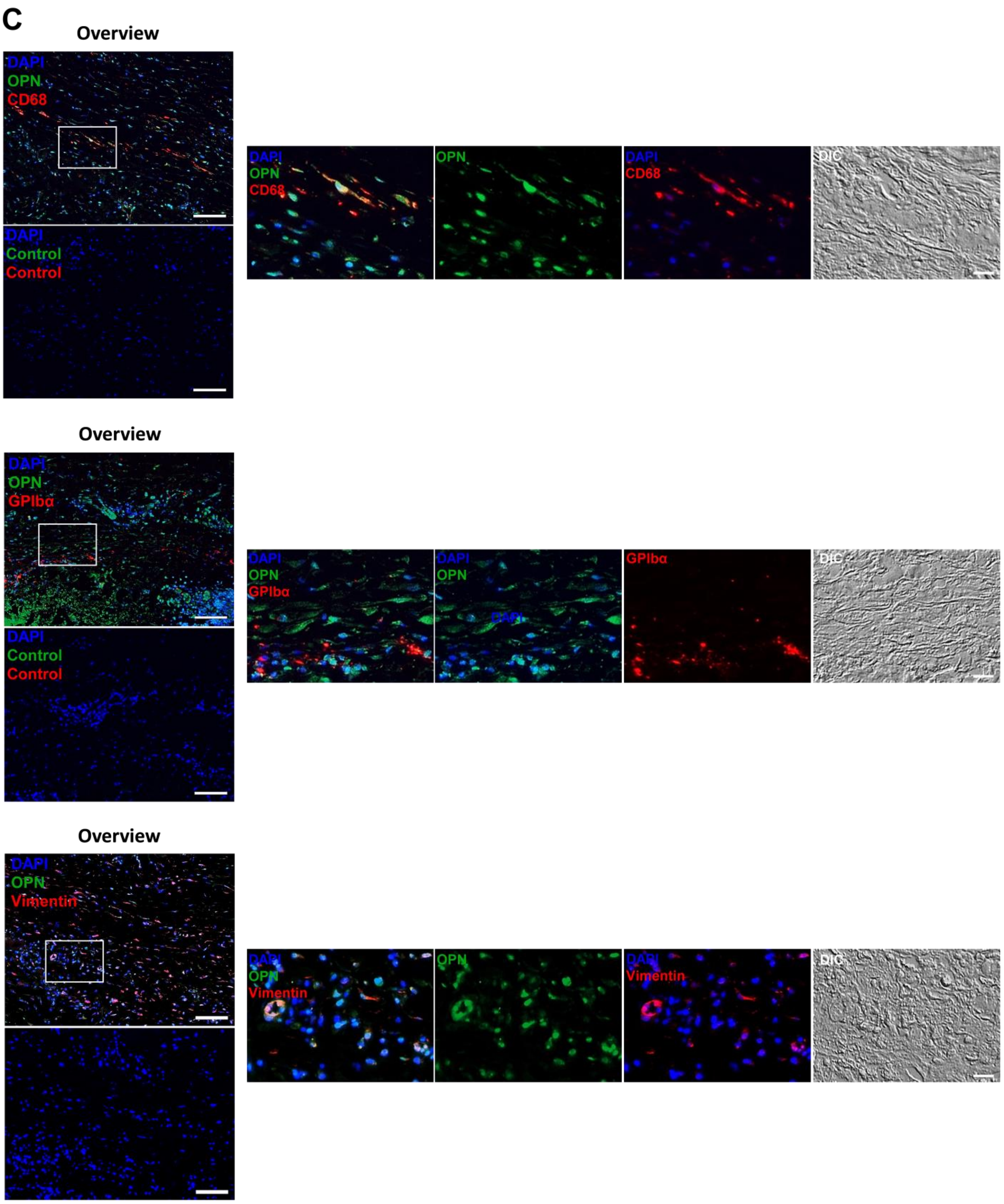


**Figure S6. Immunofluorescence staining of aortic vessel walls and intraluminal thrombi from AAA patients.** **(A+B+C)** Immunofluorescence images of **(A)** intraluminal thrombi and **(B+C)** aortic vessel wall together with negative controls were shown in an overview. Individual channels are shown in more detailed magnification. (**A**) Representative differential interference contrast (DIC) microscopy and immunofluorescence images of OPN (anti-OPN, green), platelets (CD42b/anti-GPIbα, red) cell nuclei (DAPI, blue) in specimens from the luminal layer of intraluminal thrombus (ILT) of AAA patients (n = 5). (**B**) Representative DIC and immunofluorescence images of platelets (anti-CD42b/GPIbα, 2 µg/mL, red), nuclei (DAPI, blue) and macrophages (anti-CD68 AA 21-120, 10 µg/mL, green) respectively fibroblasts (anti- vimentin, 20 µg/mL, green) in the vessel wall of AAA patients (n = 4). (**C**) Representative DIC and immunofluorescence images of osteopontin (anti-OPN, 20 µg/mL, green) and nuclei (DAPI, blue)  with macrophages (anti-CD68, 10 µg/mL, red), platelets (anti-CD42b/GPIbα, 2 µg/mL, red) and fibroblasts (anti- vimentin, 20 µg/mL, red) in aortic vessel wall (n = 4). Scale bar: 100 µm (overview) and 20 µM (detail images). Negative controls were achieved by omission of the primary antibody (control). DAPI, 4′, 6-diamidino-2-phenylindole; DIC, differential interference contrast; ILT, intraluminal thrombus; OPN, osteopontin.
